# Minimal activation of the p53 DNA damage response by a modular cytosine base editor enables effective multiplexed gene knockout in induced pluripotent stem cells

**DOI:** 10.1101/2025.06.05.656583

**Authors:** Robert Blassberg, Olga Mielczarek, Jesse Stombaugh, Hanneke Okkenhaug, Alice Abreu Torres, Paul Russell, John Lambourne, Immacolata Porreca

**Affiliations:** Cell & Gene Therapy R&D, Revvity Discovery, Cambridge, CB25 9TL, UK; Bioinformatics, Revvity, 2650 Crescent Drive, Lafayette, CO 80026, US; Imaging Facility, Babraham Institute, Cambridge, CB22 3AT, UK

## Abstract

Precise genome editing of induced pluripotent stem cells (iPSC) holds great promise for engineering advanced cell therapies. CRISPR-Cas systems have been widely adopted in genome engineering applications, however their dependence on genotoxic DNA double strand breaks (DSBs) presents challenges in hypersensitive iPSCs. Base editors are capable of both modifying and ablating gene function without generating DSBs making them an attractive solution for iPSC engineering. Here we report efficient and durable multiplexed target knockout with minimal impact on cell viability and expansion with a cytosine base editor composed of nCas9-UGI and Rat APOBEC1 assembled using the Pin-point™ platform (nCas9-UGI:rAPO). Minimal p53-mediated DNA damage signalling occurred independently of the number of simultaneous edits installed, and this could be further reduced by modulating the assembly of the base editor complex. Whereas non-homologous end-joining-mediated DNA damage repair led to p53-mediated selection against imprecise editing outcomes and an associated reduction in the on-target efficiency of multiplexed SpCas9 nuclease editing, p53 activity was dispensable for maintaining genome integrity during the base editing process. The Pin-point platform therefore enables the assembly of base editors optimised for high editing efficiency with substantially reduced risk of selecting for defective DNA damage responses inherent to DSB-dependent editing systems.

## INTRODUCTION

PSCs hold great promise as a source of cells and tissues for a range of adoptive and regenerative cell therapies, however their widespread clinical application will likely depend upon the use of allogeneic master cell banks with defined properties (1, 2). Immune rejection due to donor-recipient mismatch is a general obstacle faced by allogeneic cell therapies (3, 4) in addition to other cell-type specific challenges such as graft versus host disease (5) and exhaustion (6) in the case of adoptive T cell therapies. A number of strategies have been developed to address these requirements by knocking out the function of genes governing these obstructive biological processes (7–9). Genome editing with CRISPR-Cas represents a durable technology for accomplishing such aims that is making progress in clinical applications (10–12). However, due to their dependence on DSBs CRISPR-Cas systems are associated with the generation of large undesirable deletions at the DNA target site, which can lead to loss of complete chromosome arms (13–17). This propensity for generating structural variants is further compounded when multiple loci are edited simultaneously, which has been shown to promote the formation of interchromosomal translocations between DSB sites in multiple cell types (18–20). While these large-scale structural variants can be identified by karyotyping edited clones, smaller variants require more high-resolution screening methods, making the quality control process more laborious and costly.

p53 functions to maintain genome integrity by either stalling cell-cycle progression to allow DNA repair to take place or else eliminating defective genomes by triggering apoptosis (21).

Pluripotent cells that will normally constitute all tissues of the developing embryo are primed to undergo apoptosis in response to DSBs (22, 23), making them particularly sensitive to genome editing with CRISPR-Cas systems (24). p53 mutations occur spontaneously in iPSC cultures and accumulate due to the growth and survival advantage they confer upon mutant sub-clones (25). As DNA DSB dependent editing is normally suppressed by p53 mediated apoptosis, this has the undesirable consequence of selecting for the survival of edited products harbouring additional mutations in p53 and other components of the DNA damage-sensing pathway (26–28). Moreover, while it has become understood that a number of defined variants including those of p53 tend to occur reproducibly during iPSC culture (25, 29, 30), it cannot be guaranteed that additional as-of-yet undefined variants associated with impaired responses to DNA damage are not selected for following DSB-dependent genome editing. It is therefore desirable to develop new genome editing approaches that minimise the risk of selecting for edited iPSC clones with aberrant DNA damage responses.

Base editing is an emerging technology with the potential to overcome many of the obstacles presented by DSB-dependent editing with CRISPR-Cas, however to date only a limited number of studies have evaluated this technology in iPSCs (31–35). Whereas DSB-dependent gene knock-out relies on the generation of small insertions and deletions (indels) via imprecise non-homologous end joining (NHEJ) DNA damage repair at edited sites, base editing precisely installs desired alterations to target sequences via direct chemical modification of target nucleotide bases (36, 37). This is achieved by pairing a DNA targeting Cas component engineered to ablate DSB generating activity with a DNA modifying enzyme. Two prominent classes of base editors have been developed to date that catalyse transition substitutions via a deaminase enzyme that targets either cytosine or adenine nucleotides within the single-stranded R-loop DNA formed by the Cas component, referred to as cytosine base editors (CBEs) or adenine base editors (ABEs) respectively (38, 39). Both classes can ablate gene function by disrupting splice acceptor or donor sites, resulting in aberrant splicing and consequent loss of functional mRNAs (40). CBEs can additionally directly install STOP codons within ORFs, substantially increasing the number of targetable sequences and thereby increasing their versatility in gene knock-out applications (41, 42).

Base editor efficiency is substantially enhanced by utilising engineered Cas variants that generate a nick in the unedited DNA strand. This promotes the correction of the nicked strand via the mismatch repair pathway by utilising the edited base as a template, thereby converting the complementary base on the unedited strand (38). Cytosine base editor efficiency and fidelity is further enhanced by including a uracil-N-glycosylase (UNG) inhibitor (UGI) component that inhibits the base-excision repair pathway that would otherwise remove the edited base and replace it (34, 38, 43). Although base editing does not depend on the generation of DSBs to introduce the intended modification, studies in cell lines and primary haematopoietic stem cells have indicated that the base modifying, DNA nicking, and UNG inhibiting activities of both CBEs and ABEs can lead to DNA DSBs and associated indels, and elicit p53 mediated DNA damage responses (44–47). Moreover, it has been demonstrated that augmented UGI activity, UNG inhibition, and p53 inhibition can all enhance cytosine base editing in iPSCs (32–34). This raises the question of how the components of CBEs interact with the DNA damage pathways in pluripotent cells, which exhibit a high basal level of DNA damage repair and unusual cell-cycle checkpoints associated with their highly proliferative state (48–50).

To address these questions we exploited the Pin-point platform, which is a novel modular base editor system that enables assembly of independent DNA binding and modifying components via an aptamer encoded in the sequence targeting guide RNA (51, 52). We show that mRNA delivery of a CBE composed of Rat APOBEC1 and a nickase variant of SpCas9 fused to UGI (nCas9-UGI:rAPO) optimised for iPSC editing led to efficient, durable target knockout and elicited a substantially reduced p53 response and enhanced cell viability compared to SpCas9, both when a single target was edited or when multiple targets relevant to allogeneic adoptive cell therapy manufacturing were edited simultaneously. The minimal p53 pathway activity associated with the nCas9-UGI:rAPO editor was both insufficient to adversely impact multiplexed editing efficiency and dispensable for maintaining genome integrity during the base editing process, and could be further reduced by modulating assembly of the nCas9-UGI and APOBEC1 components by optimising the concentration of the aptamer encoding sgRNA. By contrast, SpCas9 editing exhibited a substantially elevated p53 response that was correlated with the number of DSBs undergoing NHEJ repair. While this p53 response acted to maintain genome stability during multiplexed editing this came at the expense of reducing on-target editing efficiency. We therefore propose that optimised formulations of Pin-point cytidine base editors enable high efficiency, high fidelity multiplexed gene knock-out applications in iPSCs likely to be necessary for engineering allogeneic master cell banks, while substantially reducing the risk of selecting for products with aberrant responses to DNA damage.

## MATERIAL AND METHODS

### Human induced pluripotent stem cell culture

A18945 (Gibco) episomal hiPSCs (53) were obtained from ThermoFisher; WTC11 hiPSCs (54) were obtained from Corriell Institute; NH50191 hiPSCs were obtained from Infinity BiologiX. All hiPSCs were cultured on Geltrex™ (ThermoFisher) matrix coated cell culture plastic in mTesr™ PLUS defined medium (STEMCELL Technologies) at 37°C, 5% CO_2_. Sub-confluent cultures were passaged by non-enzymatic dissociation with Versene™ solution (ThermoFisher) and re-plated as clumps.

### Guide RNA design

All gene-targeting gRNAs used in this study were identified and characterised previously (52). Briefly, a combination of manual and automated sequence inspection was employed to identify base conversions at intron-exon boundaries that would disrupt splice donors or acceptors. Guides that targeted more than a single location within the genome were removed from consideration. The non-targeting control guide (Revvity) was bioinformatically optimised to minimise homology with off-target sequences. sgRNAs used with the modular Pin-point base editing system included the MS2 phage operator stem-loop aptamer while sgRNA used with SpCas9 did not include the aptamer. Details of base editing guide RNAs are reported in Supplementary Table 1.

### Editing reagents

HPLC purified sgRNAs were synthesized by Agilent Technologies or Revvity and diluted in Tris Buffer. mRNAs encoding components of the Pin-point system (nCas9-UGI, dCas9-UGI, nCas9, and rAPOBEC1-MCP) and SpCas9 were fully substituted with Pseudo-Uridine and 5-methyl-cytosine and capped with CleanCap™ mRNA capping technology (TriLink BioTechnologies). nCas9-UGI was derived from SpCas9 D10A, and dCas9-UGI was derived from SpCas9 D10A H840A (55). mRNA sequences are available in Supplementary table 2.

### hiPSCs Electroporation

3hr prior to electroporation sub-confluent iPSC cultures were treated with 10uM Rho kinase inhibitor Y-27632 (Stemcell Technologies). hiPSC colonies were dissociated to single cells using StemPro™ Accutase™ cell dissociation reagent (ThermoFisher) prior to resuspension in electroporation buffer P3 (Lonza). Dissociated iPSC samples (2.5 x 10^5^ per electroporation) were electroporated with 40 pmol of each gRNA and either 1.6 pmol (2.5 ug) of the nCas9-UGI and 1.6 pmol (0.75 ug) of the APOBEC1-MCP base editor components, or 1.5 pmol (2 ug) of SpCas9 unless otherwise indicated using the 20ul Nucleocuvette™ strip Amaxa™ 4D Nucleofector® System (Lonza). After electroporation, hiPSCs were transferred directly to Geltrex coated cell culture vessels containing prewarmed mTesr-PLUS supplemented with 10uM Rho kinase inhibitor Y-27632 and incubated at 37°C, 5% CO_2_ for 24hrs. Culture medium was then changed for mTesr-PLUS without additional Y-27632 and electroporated hiPSCs were expanded with daily complete medium changes.

For p53 knockdown experiments, 20 pmols of Dharmacon™ ON-TARGETplus™ siRNA SMARTpool™(Revvity) diluted in Tris buffer targeting either p53 or a non-targeting control pool were included with editing reagents in the electroporation.

### Small molecule treatment

Where used, the MDM2 inhibitor Nutlin-3a (Cayman Chemical) was included in culture medium at the indicated concentration during the 24 hours following electroporation of editing reagents.

### Genotyping of genomic DNA samples

For gDNA preparation, hiPSCs were lysed using DirectPCR® cell lysis buffer (Viagen Biotech, LA, USA) supplemented with proteinase K (10 µg/mL; Sigma-Aldrich) and heated at 55°C for 60 min, then 95°C for 20 min. The crude lysate was then used as input material for PCR using locus-specific primers with universal adapters designed to amplify a 250-350 bp site surrounding the genomic region of interest (for primer sequences see Supplementary Table 3). PCR products were cleaned up and sequenced by Sanger sequencing using a universal sequencing primer (Genewiz, Azenta). For quantification of base conversion, AB1 files were analysed using custom software based on the BEAT tool (56). For quantification of indels conversion AB1 files were analysed using custom software based on the TIDE tool (57).

### Flow cytometry

For labelling of cell surface antigens, cells were dissociated with StemPro Accutase cell dissociation reagent (ThermoFisher) and incubated for 15 mins at 4oC with fluorophore-conjugated antibody diluted in PBS [human B2M (BioLegend, #316312), 1:1000]. Labelled cells were pelleted at 400*g* for 5 min, unbound antibody was removed, and samples were washed by resuspension in PBS. Cell viability was assessed using DAPI (BD, #564907; 80 ng/mL). For intracellular staining, dissociated cells were pelleted and resuspended in 4% paraformaldehyde in PBS. Following 15 min incubation at 4 °C, cells were centrifuged at 700 *g*, resuspended in PBS and stored at 4 °C for future analysis. For staining samples were pelleted in V-bottom 96-well plates, resuspended in FACS block (PBS + 0.2% Triton + 3% BSA), and incubated at room temperature for 10 mins. Antibody was added to the sample and incubated overnight at 4 °C [human OCT3/4 (BD #560307), 1:200]. Labelled cells were pelleted at 700*g* for 5 min and resuspended in 50 μl FACS block. One additional wash was performed before acquisition.

Acquisition was performed on an IntelliCyte™ IQue PLUS or Sartorius iQue3® flow cytometer using iQue ForeCyt® Enterprise Client Edition 9.0 (R3) Software for both acquisition and data analysis.

### Live-imaging of cell growth (Incucyte)

Electroporated cells were plated on Geltrex matrix coated 96 well plates (Corning) in mTesr-PLUS (STEMCELL Technologies) medium supplemented with 10uM Rho kinase inhibitor Y-27632 and cultured in an Incucyte® imager housed in an incubator set at 37°C, 5% CO_2_.

Following 24hrs in culture, medium was changed for fresh mTesr-PLUS (STEMCELL Technologies), and every 48 hrs thereafter. Cell confluence was determined using the Incucyte 2021C software.

### Cell cycle analysis and apoptosis assays

Electroporated cells were plated on Geltrex matrix coated 96 well plates (Corning) in mTesr-PLUS (STEMCELL Technologies) medium supplemented with 10uM Rho kinase inhibitor Y-27632 and incubated at 37°C, 5% CO_2_. Following the indicated period in culture cells were cultured for a further 4 hrs in the presence of 5ug/ml Hoechst 33342, dissociated to single cells using StemPro Accutase cell dissociation reagent (ThermoFisher), and equivalent proportions of each sample were labelled for 20 minutes at room temperature with Alexa Fluor®-488 conjugated Annexin V (ThermoFisher) diluted 1:40 in Annexin V Binding Buffer (10mM Hepes pH7.4, 140mM NaCl, 2.5mM CaCl2), and Propidium Iodide (ThermoFisher) in 96 well plates using a minaturised version of the manufacturers protocol. Cells were then labelled with Propidium Iodide for 15 mins at room temperature. Acquisition was performed on an IntelliCyte IQue PLUS or Sartorius iQue3 flow cytometer using iQue ForeCyt Enterprise Client Edition 9.0 (R3) Software for both acquisition and data analysis.

The proportions of live, early apoptotic, late apoptotic and dead cells were determined from Annexin V and Propidium Iodide co-staining, setting gating thresholds on the mock electroporated sample. Viable cell counts were determined from the propidium iodide negative population. Cell cycle phase (DNA content) was determined from the linear Hoechst 33342 fluorescence area signal. Representative graphs of DNA content distributions and Annexin V/Propidium Iodide co-labelling were generated using the R packages flowCore and ggplot2.

### High-content imaging

Electroporated iPSCs were seeded onto Geltrex (ThermoFisher) matrix coated 96-well PhenoPlate™ microplates (Revvity) in mTesr-PLUS (STEMCELL Technologies) supplemented with 10uM Rho kinase inhibitor Y-27632 (STEMCELL Technologies) and incubated at 37°C, 5% CO_2_. Cells were fixed with 4% paraformaldehyde in PBS for 15 minutes at 4oC at the indicated time point then rinsed twice with PBS. Samples were blocked and permeabilised in Immuno Block solution (PBS + 0.1% Triton + 1% BSA) for 10 min at room temperature, after which primary antibodies were added at the concentrations indicated in Supplementary Table 4 in fresh Immuno Block solution, and incubated overnight at 4oC. Primary antibody was removed and samples were washed with Immuno Block solution 3 times for 5 minutes, after which secondary antibodies plus 1ug/ml Hoechst 33342 stain (ThermoFisher) were added at the concentrations indicated in Supplementary Table 4 in fresh Immuno Block solution, and incubated for 2 hrs at room temperature. Secondary antibody was removed and samples were washed with Immuno Block solution 3 times for 5 minutes, followed by rinsing with PBS and mounting in PBS for imaging.

Imaging was performed using an Opera Phenix^™^ Plus high-content screening system (Revvity) through a 40x 1.1NA water immersion lens, or at the Babraham Institute Imaging Facility using an ImageXpress® Confocal HT-AI High Content Imaging System (Molecular Devices) through a 20x 1.2NA water immersion lens.

### Image segmentation and analysis

Image segmentation was performed in Harmony™ software (Revvity) or at the Babraham Institute Imaging Facility using CellProfiler™ 4.2.1 software (Broad Institute) to identify and quantify the following features: Individual Hoechst stained nuclei regions of interest (NucROI); Average p53 pixel intensity within individual NucROIs (Mean_p53); Area of NucROIs; Individual γH2AX DNA repair foci within individual NucROIs (γH2AX_ROI); Area of γH2AX_ROIs; Total 53BP1 pixel intensity within H2AXy_ROIs; Individual 53BP1 foci within individual NucROIs (53BP1_ROI); Area of 53BP1_ROIs; Count of Hoechst stained micronuclei associated to Nuclei.

Custom R scripts were created to calculate metrics presented in figures. Percentage of p53_high nuclei in each sample was calculated by first calculating the Mean_p53 for each NucROI (total p53 pixel intensity within a given NucROI/ Area of given NucROI), then determining the percentage of NucROIs with Mean_p53 greater than a threshold set in relation to the mock electroporated samples within an experiment. Average number of small and large foci per nucleus were calculated by first categorising individual γH2AX_ROIs or 53BP1_ROIs (foci_ROIs) as large or small based on area, with the threshold set in relation to the mock electroporated samples within an experiment, then dividing the total number of large and small foci_ROIs by the number of NucROIs within a given sample. The probabilities of a nucleus having high levels of p53 given the number of small foci_ROIs (n_foci_small_) or large foci_ROIs (n_foci_large_), P(p53_high|n_foci_small_) and P(p53_high|n_foci_large_) respectively, was determined by first binning NucROIs by n_foci_large_ and then calculating the percentage of p53_high NucROIs within each bin as described above; NucROIs with n_foci_large_ = 0 were then binned by n_foci_small_ and the percentage of p53_high NucROIs within each bin was calculated by the same method.

Representative distributions of Mean_p53 levels, γH2AX_ROI area and 53BP1_ROI area distributions were generated with ggplot.

### cDNA synthesis and qPCR analysis

First-strand cDNA synthesis was performed using Superscript™ III reverse transcriptase (Invitrogen) using random hexamers and was amplified using PowerUp™ SYBR™ Green Mastermix (Applied Biosystems). qPCR was performed using the QuantStudio™ 6 Flex Real Time PCR system (Applied Biosystems) and analysed with QuantStudio software (Applied Biosystems, V1.7.2). PCR primers were designed using online qPCR primer design tool (GenScript). Two technical replicates were obtained for each sample and averaged before normalization and statistical analysis. Relative expression values for p21 were calculated by normalization against GAPDH, using the delta–delta CT method. qPCR analysis was performed on samples obtained from a minimum of 2 independent experiments. Data were graphed and statistical tests were performed using GraphPad Prism software. Primer sequences are detailed in Supplementary Table 5.

### Capture-seq

Hybridisation capture sequencing was performed as described (52) with the inclusion of additional probes designed to enrich guide RNA off-target sites identified in the previous study (for sequences see Supplementary Table 6).

*Library preparation.* Briefly, genomic DNA samples were prepared from sub-confluent edited iPSC cultures using DNeasy® Blood and Tissue Kit (Qiagen). 1µg of gDNA were used to prepare paired-end sequencing libraries using the KAPA HyperPlus™ DNA library preparation kit (Roche) followed by enrichment of genomic regions flanking gene editing targets using the xGen™ NGS Hybridization Capture workflow with xGen™ Hybridization Capture Core Reagents (IDT) as described previously (52). Libraries were sequenced on an Illumina MiSeq™ sequencing system in 2×300bp runs (Illumina, San Diego) by Source Bioscience.

*Read alignment and structural variant identification.* Sequencing reads were trimmed to the first 75bps using a custom Python script and then processed through the DRAGEN™ Structural Variant (SV) Caller (58) (version 3.8.4) (Illumina) to identify structural variants, which extends the MANTA (59) structural variation pipeline. For detection of translocations, samples were run through the pipeline as an unpaired tumor samples. For detection of insertions and deletions, samples were run through the pipeline as tumor-normal pairs using corresponding mock edited parental line samples as normal. An example command is as follows:

/opt/edico/bin/dragen-f--ref-dir /ephemeral/ucsc.hg38.3.8.4/--tumor-fastq1

s3://aws_bucket/Sample1_R1_001.paired.75bp.fastq.gz--tumor-fastq2

s3://aws_bucket/Sample1_R2_001.paired.75bp.fastq.gz--output-directory

/ephemeral/DRAGEN_Sample1/--output-file-prefix Sample1--enable-duplicate-marking

true--enable-map-align true--enable-map-align-output true--enable-sv true--RGID-tumor Sample1--RGSM-tumor Sample1--sv-exome true--remove-duplicates true

*Translocations quantification.* To remove reads derived from library fragments captured non-specifically during hybridisation the *“*.candidateSV.vcf*” output from MANTA was first filtered to include only breakends within sequences mapping to genomic regions +/-1000bp either side of gene editing targets. Interchromosomal translocations were quantified by normalising the total count of reads (BND_PAIR_COUNT) supporting a given variant involving regions on two different chromosomes by the total number of reads mapping to either genomic region fusion point on each chromosome. Where both genomic regions adjacent to the breakpoint contained sequences targeted by capture probes the average number of reads across these regions was used for normalisation.

## RESULTS

### Minimal p53 pathway activation by the optimised nCas9-UGI:rAPO base editor

To investigate the utility of the Pin-point base-editing platform for gene knockout in iPSCs we employed synthetic Pin-point aptamer-encoding sgRNAs and mRNA reagents previously optimised for the generation of allogeneic CAR-T cells (52) (Fig 1A). We exploited the modularity of the system to define the optimal ratio of the sgRNA, nCas9-UGI, and Rat APOBEC1-MCP (rAPO) modules for maximal editing (Fig S1A) while minimising cytotoxicity (FigS1 B,C). This was determined to be a 1:1 molar ratio of nCas9-UGI:rAPO mRNAs and 40 pmol of sgRNA, and was used in all experiments described in the study unless otherwise stated.

**Figure 1:**
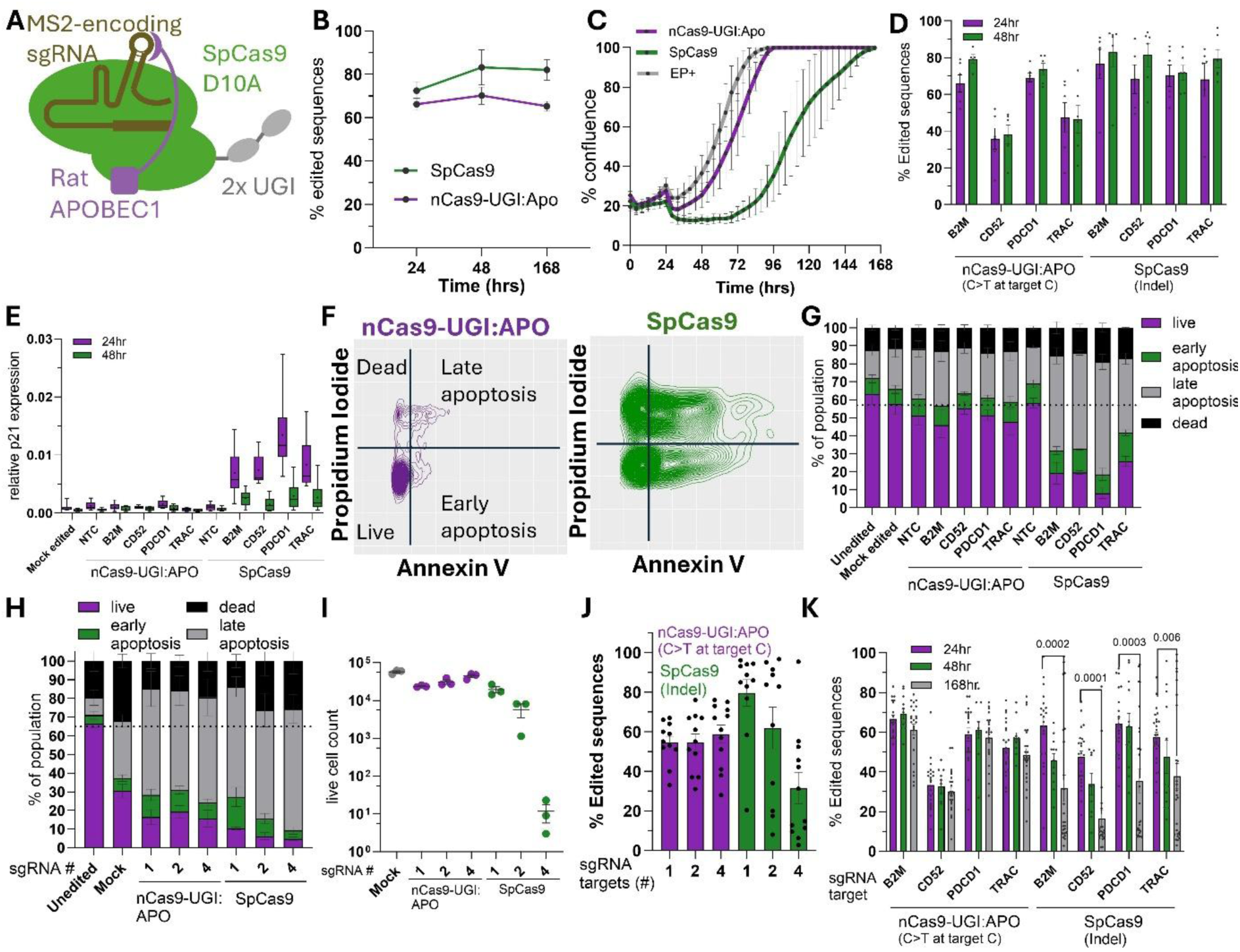
Minimal p53 mediated apoptosis by the optimised nCas9-UGI:rAPO base editor. A) Schematic of the components of the modular Pin-point cytidine base editor. B) Percentage of either SpCas9-induced indels or nCas9-UGI:APO-induced base conversion at the target nucleotide over time following delivery of editors targeting *B2M* analysed by Sanger sequencing. Mean +/-SEM, n=3 iPSC lines. C) Confluence of cultures over time following delivery of the editor analysed by time-lapse microscopy. Mean +/-SEM, n=3. D) Percentage of either SpCas9-induced indels or nCas9-UGI:APO-induced base conversion at the target nucleotide over time following delivery of editors targeting the indicated genes analysed by Sanger sequencing. Mean +/-SEM. E) p21 expression over time following delivery of editors targeting the indicated genes. n=6-12. Tukey plots with mean shown by *. F) Representative flow-cytometry plots of Annexin V apoptosis assay 48 hours following delivery of editors targeting the *PDCD1* gene. G) Quantification of Annexin V apoptosis assay 48hrs following delivery of editors targeting the indicated genes. Mean +/-SEM, n=3. H) Quantification of Annexin V apoptosis assay 48 hours following delivery of editors targeting the indicated number of genes simultaneously. Mean +/-SEM, n=3-6. I) Quantification of propidium iodide negative live cells assayed 7 days following delivery of editors targeting the indicated number of genes simultaneously. Mean +/-SEM. J) Percentage of either SpCas9-induced indels or nCas9-UGI:APO-induced base conversions at the target nucleotide within the *B2M* locus following delivery of editors targeting the indicated number of genes simultaneously. Analysed by Sanger sequencing 7 days following delivery of editors. Mean +/-SEM. K) Percentage of either SpCas9-induced indels or nCas9-UGI:APO-induced base conversion at the target nucleotide over time following delivery of editors targeting four genes simultaneously analysed by Sanger sequencing. Mean +/-SEM. p-values from paired t-tests.

To investigate the long-term impact of base-editing on iPSC fitness we subcultured pools of edited cells for 100 days and monitored expression of the target protein B2M and the pluripotency marker OCT4. Pluripotency was stably maintained following base editing (Fig S1D) and the percentage of B2M negative edited cells remained constant in subcultured pools (Fig S1E), both confirming the durability of the DNA base edit and indicating that edited pluripotent cells exhibited neither a selective disadvantage nor advantage over unedited cells.

We next sought to benchmark viable cell expansion of iPSCs edited with nCas9-UGI:rAPO against DSB-based editing using an equal molar adjusted quantity of SpCas9 mRNA targeted to the same genomic locus with an sgRNA lacking the aptamer sequence. We first defined the editing kinetics of each system, which demonstrated that both base editing by nCas9-UGI:rAPO and indel formation by SpCas9 occur rapidly, with the majority of editing occurring within the first 24hrs following delivery of the RNA components of each system (Fig 1B).

Time course imaging analysis of the kinetics of cell expansion following editing demonstrated that iPSCs edited with nCas9-UGI:rAPO began to expand exponentially from 24hrs once recovered from the initial stress of electroporation observed in mock edited controls. By contrast, Cas9 edited iPSCs did not begin to expand for a further 48hrs (Fig 1C), indicating that expression of the base editing machinery is markedly less deleterious to iPSC expansion than SpCas9.

As the p53 pathway has been shown to trigger apoptosis of SpCas9 edited iPSCs (24) we hypothesised that p53 activity may explain the differences in iPSC expansion associated with the two editing systems. To test this hypothesis, we expanded our analysis to a set of sgRNAs previously optimised for functional knock-out in primary T-cells (52). These sgRNAs exhibited a broad range of editing efficiencies in iPSCs (Fig 1D), providing the opportunity to investigate the relationship between efficiency, p53 activation, and cytotoxicity for both editing systems. When used to target SpCas9, all sgRNAs showed a transient activation of the p53 pathway compared to mock edited controls. By contrast, when used to target the nCas9-UGI:rAPO base editor even the most efficiently editing sgRNAs showed no greater p53 pathway activation than either a non-targeting sgRNA or the mock edited control (Fig 1E). Consistent with the observation that the p53 pathway was activated exclusively in Cas9 edited iPSCs, we observed an increase in the late apoptotic fraction when SpCas9 was targeted by any of the four sgRNAs (Fig 1F,G). These effects were most apparent for the sgRNA targeting PDCD1 which we have shown previously to have a high efficiency off-target site (52), and therefore targets multiple sites in the genome. By contrast, none of the sgRNAs impacted cell viability when used to target nCas9-UGI:rAPO (Fig 1F,G).

### Durable high efficiency multiplexed editing with the nCas9-UGI:rAPO base editor

It has been shown previously that targeting multiple sites in the genome simultaneously with Cas9 amplifies p53 mediated effects (60). We therefore investigated whether base editing with nCas9-UGI:rAPO further enhanced the recovery of viable edited iPSCs compared to SpCas9 when multiple loci were targeted simultaneously. To control for increased quantities of sgRNA used for multiplexed editing we delivered the same 40 pmol quantity of B2M sgRNA shown to achieve the optimal balance of editing and cell viability in combination with additional targeting sgRNAs while keeping the total sgRNA quantity constant (160 pmol) with non-targeting sgRNA. Increased numbers of SpCas9 edits were associated with an initial increase in the proportion of dead and apoptotic cells and a dramatic reduction in viable cell counts following multiplexed editing of four targets and 7 days of subsequent expansion (Fig 1H,I). By contrast, the proportion of apoptotic cells was unaffected by simultaneous editing of 4 targets with the nCas9-UGI:rAPO base editor, with the yield of viable cells remaining equivalent to the mock edited control (Fig 1H,I). Similarly, simultaneously editing of multiple targets with SpCas9 led to a drop in SpCas9 editing efficiency compared to when only a single target was edited (Fig 1J), whereas the editing efficiency of the nCas9-UGI:rAPO base editor was unaffected by multiplexed editing.

The distinct relationships between cell viability and editing efficiency observed when simultaneously editing multiple targets with the two technologies indicated that apoptosis specifically selects against multiplexed SpCas9 editing. We reasoned that if this was the case, the frequency of edited cells would be observed to decrease over time in SpCas9 but not nCas9-UGI:APO edited populations. Consistent with this hypothesis, simultaneously targeting multiple loci with SpCas9 led to a drop in mean editing efficiency over time for the four targets in each of the cell lines analysed, whereas the frequency of nCas9-UGI:rAPO base edited sequences remained constant (Fig 1K, S1F-H). Taken together these data demonstrate that base editing with nCas9-UGI:rAPO avoids selection against edited iPSCs that occurs when multiple targets are edited simultaneously by SpCas9.

### On-target editing with nCas9-UGI:APO is unaffected by low level p53 pathway activity

Having previously observed that single target editing with SpCas9 led to substantially greater p53 pathway activation than the nCas-UGI:rAPO editor (Fig 1E) we investigated whether the favourable properties of the nCas-UGI:rAPO editor in multiplexed editing applications could also be explained in terms of p53 pathway activation. Indeed, whereas simultaneous editing of 4 targets with SpCas9 resulted in a substantially greater increase in nuclear p53 levels (Fig 2 A,B) and p21 induction (Fig 2C) than editing of 1 target, editing of 4 targets with nCas9-UGI:rAPO base editor resulted in only a moderate increase in p53 activity compared to single target editing (Fig2 A-C). These results indicated that minimal p53 induced apoptosis likely explains the enhanced multi target editing efficiency associated with the nCas9-UGI:rAPO base editor.

**Figure 2:**
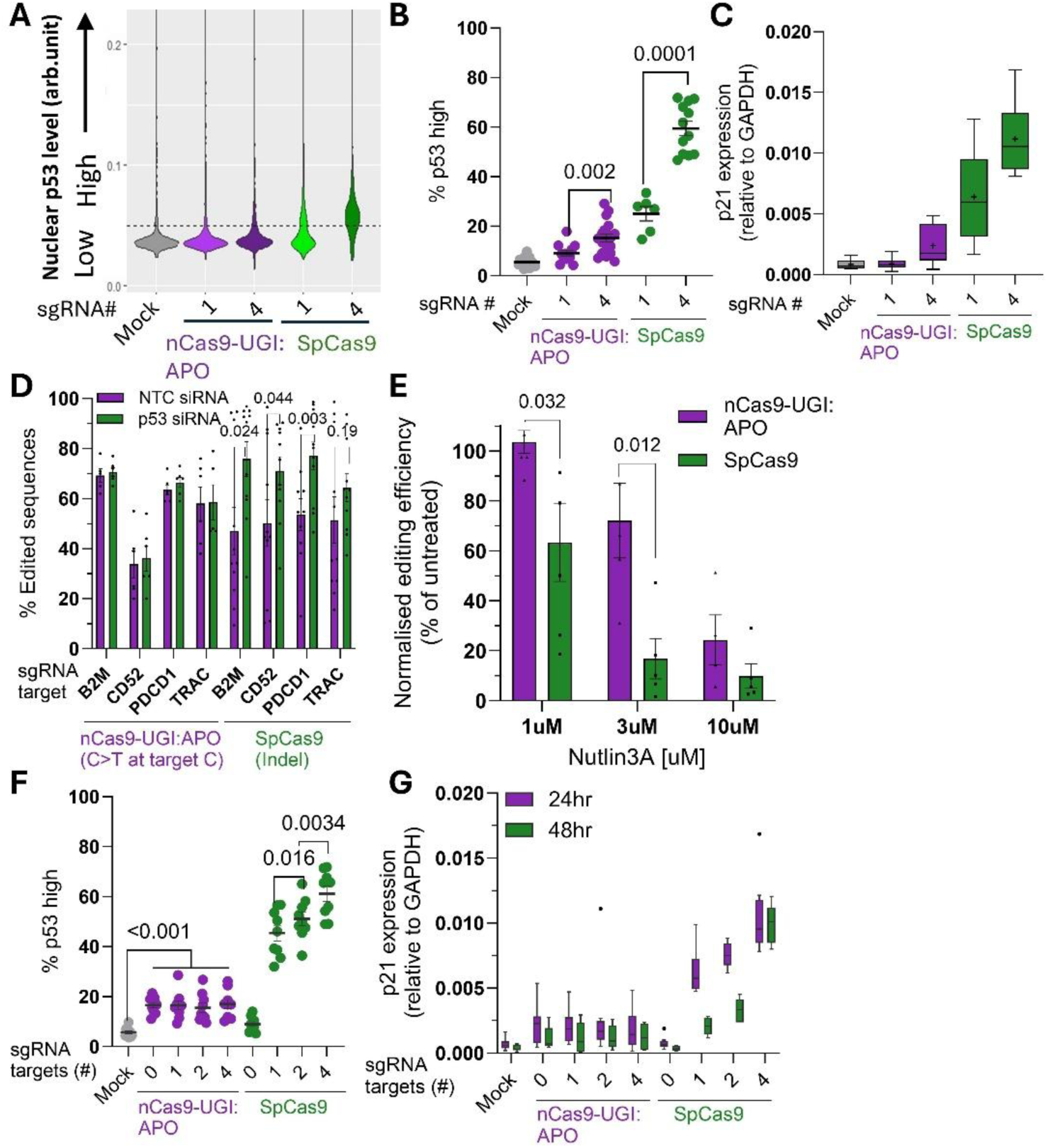
On-target editing with nCas9-UGI:APO is unaffected by low level p53 pathway activity. A) Representative distributions of average nuclear p53 levels following single target or multiplexed editing. p53 levels above the main distribution of mock edited controls were classified as ‘high’. B) Quantification of the percentage of nuclei classified as containing high levels of p53 by microscopy in mock edited samples, or samples edited with the nCas9-UGI:APO editor or SpCas9 at either 1 or 4 targets simultaneously, analysed 24 hours following deliver of the editor. Mean +/-SEM. C) Expression of the p53 pathway target gene p21 in mock edited samples, or samples edited with either the nCas9-UGI:APO editor or SpCas9 at either 1 or 4 targets simultaneously. Analysed 24hrs following deliver of the editor.

As it has been reported that p53 activity inhibits efficient base editing (33) we directly tested whether the low level of p53 activity associated with multiplexed editing with nCas9-UGI:rAPO impacted the editing process. Consistent with previous reports (61), transient inhibition of p53 pathway activity by depletion of nuclear p53 with siRNA (Fig S2 A-C) enhanced editing efficiency of SpCas9 multi target edited iPSCs (Fig 2D). By contrast, this was not observed in the case of nCas9-UGI:rAPO (Fig 2D). This suggested to us that the low levels of p53 pathway activity elicited by multiplexed editing with the optimised nCas9:APO editor formulation were insufficient to select against edited cells. To directly test this, we increased p53 activity by stabilising nuclear p53 levels using the MDM2 inhibitor Nutlin3A (Fig S2 D-G). Consistent with this hypothesis we observed a correlated reduction in editing efficiency and cell viability for both nCas9-UGI:rAPO and SpCas9 (Fig 2E, S2 H). However, consistent with the reduced levels of p53 activity elicited by nCas9-UGI:rAPO, editing efficiency was significantly less sensitive to MDM2 inhibition, requiring higher concentrations to reach comparable levels of nuclear p53 and associated reduction in editing efficiency observed for SpCas9 (Fig 2E, Fig S2 E,F). Taking these data together we therefore conclude that at sufficiently high levels of p53 activity nCas9-UGI:APO base editing efficiency can be reduced due to p53 mediated cell apoptosis, consistent with previous reports describing other cytidine base editors (32, 33). However, as this level is not reached even when multiple loci are targeted simultaneously by transient RNA delivery of the nCas9-UGI:rAPO editor, editing efficiency remains unaffected.

Although the levels of p53 pathway activity associated with nCas9-UGI:rAPO were insufficient to inhibit editing efficiency we nonetheless sought to understand why p53 activity was elevated by multiplexed editing to further improve the safety profile of the editor. To distinguish whether p53 activity depended upon the number of sites targeted simultaneously or merely the quantity of sgRNA used in the editing process we again maintained the total quantity of sgRNA constant with a non-targeting control while increasing the number of sites targeted by either SpCas9 or nCas9-UGI:rAPO. This revealed that whereas the number of loci targeted simultaneously with Cas9 was correlated with both an initial increase in nuclei with high levels of p53 (Fig 2F, S3 A) and an extended duration of p53 target gene expression (Fig 2G, S3 B-D), nuclear p53 and downstream target gene expression was moderately increased above the levels of mock electroporated controls irrespective of the number of loci targeted by the nCas9-UGI:rAPO base editor.

Taken together the data indicated a potentially editing-independent effect of sgRNA quantity on p53 pathway activation by the nCas9-UGI:rAPO editor. Consistent with this hypothesis, p53 activity was correlated with the quantity of non-targeting sgRNA delivered in combination with the nCas9-UGI:rAPO editor (Fig S3 E,F). Thus, while the low level p53 pathway activation associated with multiplexed base editing occurred independently of on-target editing activity, multiplexed editing with SpCas9 lead to increasing p53 pathway activity proportional to the number of targets simultaneously edited.

Tukey plots with mean shown by *. D) Percentage of either SpCas9-induced indels or nCas9-UGI:APO-induced base conversion at the target nucleotide following delivery of editors targeting four genes simultaneously in combination with either a non-targeting (NTC) or p53 targeting siRNA, analysed by sanger sequencing 7 days following delivery of editors. Mean +/-SEM, p-values from paired t-tests. E) Percentage of either SpCas9-induced indels or nCas9-UGI:APO-induced base conversion at the target nucleotide following delivery of editors targeting four genes simultaneously in combination with the indicated concentration of the p53-stabilising compound Nutlin3A. Editing efficiency was analysed by Sanger sequencing 7 days following delivery of editors and is expressed relative to matched untreated control sample. Mean +/-SEM. p-values from paired t-tests. F) Quantification of the percentage of nuclei classified as containing high levels of p53 by microscopy in mock edited samples, or samples edited simultaneously at the indicated number of targets, analysed 24 hours following delivery of the editor. Mean +/-SEM, p-values from paired t-tests. G) p21 expression in mock edited samples, or samples edited simultaneously at the indicated number of targets, analysed at the indicated timepoints following delivery of the editor. Tukey plots with mean shown by *.

### p53 activity is not required to maintain genome integrity during multiplexed editing with the nCas9-UGI:rAPO base editor

As the activation of p53 signalling by the nCas9-UGI:rAPO editor occurred independently of on-target editing, this presented an opportunity to further reduce the p53 response while maintaining multiplexed editing efficiency. It has been proposed that base editing activates p53 signalling due to the incidental generation of DNA DSBs (47), motivating us to further understand the relationship between p53 activation and DNA damage associated with editing by quantifying markers of DNA DSB repair and p53 levels in individual nuclei. Consistent with previous reports that cycling iPSCs undergo DNA double strand breaks due to replication stress (49), mock edited control populations had within them a proportion of nuclei with detectable DNA repair foci marked by the general DSB marker phospho-H2AX (γH2AX) (62) (Fig 3A,B S4A). However, these foci were notably smaller than those detectable in SpCas9 edited samples, suggestive of a qualitative difference in these two classes of DNA breaks (Fig 3B, S4A). Consistent with previous reports from other cell types that base editors can cause DNA DSBs (44, 46, 47), nCas9-UGI:rAPO editing also resulted in an increase in γH2AX DNA repair foci compared to unedited controls, with this increase being more apparent when multiple loci were targeted simultaneously (Fig 3B S4A). However, whereas SpCas9 editing was predominantly associated with an increase in large foci, an increase in small γH2AX foci was more pronounced in nCas9-UGI:rAPO edited samples (Fig 3B S4A). As observed with p53 pathway activation, the frequency of γH2AX foci generated by nCas9-UGI:rAPO was also correlated with the quantity of non-targeting sgRNA delivered (Fig S4B), indicating that the sgRNA dose dependent effect on p53 activity is likely due to untargeted DNA damage at higher sgRNA quantities.

**Figure 3:**
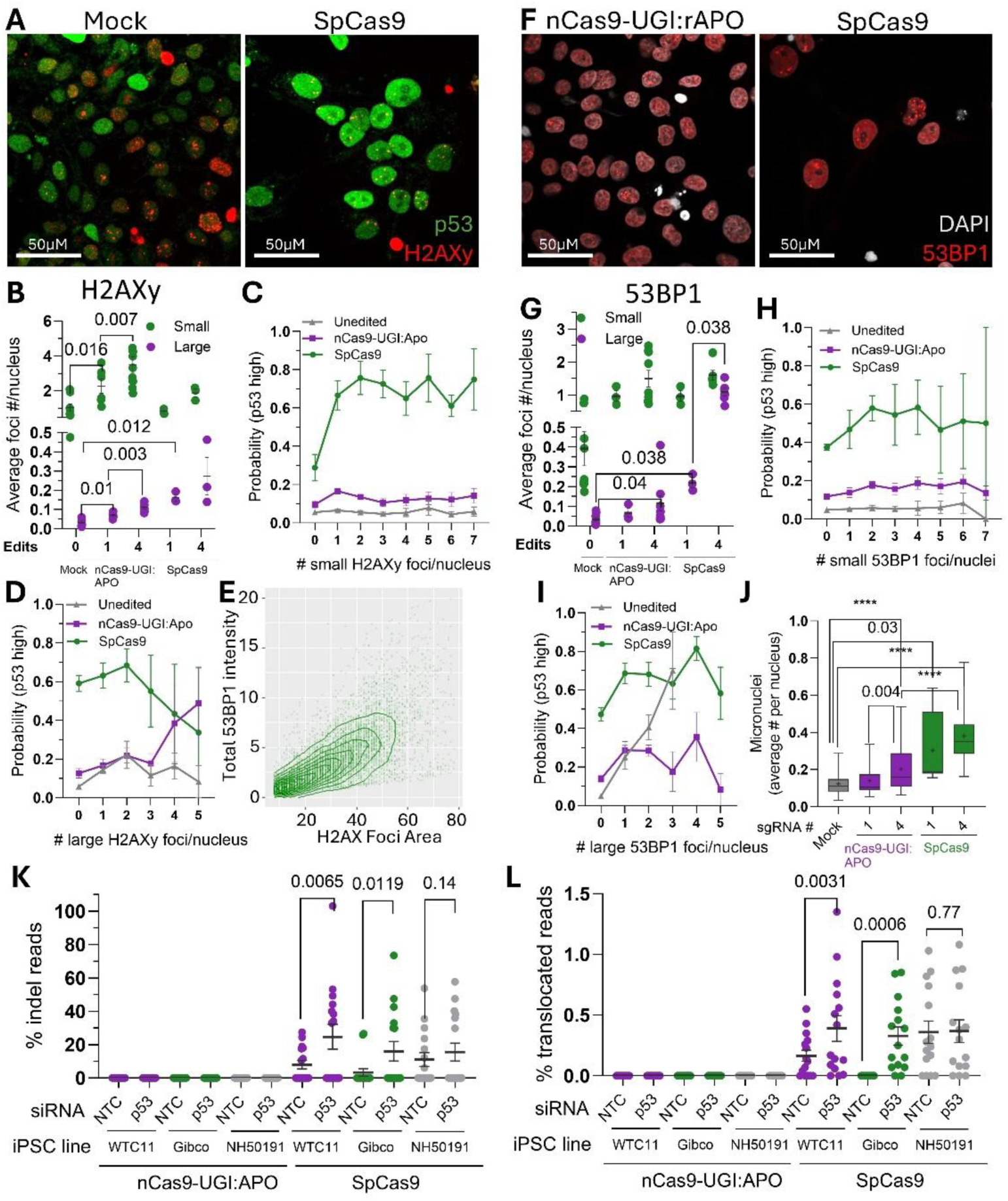
p53 activity is not required to maintain genome integrity during multiplexed editing with the nCas9-UGI:rAPO base editor. A) Representative microscopy image used for simultaneous quantification of p53 levels and DNA repair foci in single nuclei. B) Quantification of the average number of γH2AX foci per nucleus classified as either large or small 24hrs following delivery of editors targeting the indicated number of genes simultaneously. Mean +/-SEM. Quantification of the probability of nuclei containing high levels of p53 by microscopy in populations of nuclei containing the indicated number of either C) small, or D) large γH2AX foci. Nuclei in C) contain 0 large foci, whereas nuclei in D) contain an undefined number of small foci. Analysed 24hrs following delivery of editors targeting 4 genes simultaneously. Mean +/-SEM. E) Correlation of 53BP1 levels with γH2AX foci size in individual foci present in nuclei 24hrs following delivery of the SpCas9 editor targeting 4 genes simultaneously. F) Representative image of 53BP1 foci 24hrs following delivery of editors targeting 4 genes simultaneously. G) Quantification of the average number of 53BP1 foci per nucleus classified as either large or small 24hrs following delivery of editors targeting the indicated number of genes simultaneously. Mean +/-SEM. Quantification of the percentage of nuclei containing high levels of p53 by microscopy in populations of nuclei containing the indicated number of either H) small, or I) large 53BP1 foci. Nuclei in H) contain 0 large foci, whereas nuclei in I) contain an undefined number of small foci. Analysed 24hrs following delivery of editors targeting 4 genes simultaneously. Mean +/-SEM. J) Quantification of the average number of micronuclei associated with nuclei 24hrs following delivery of editors targeting the indicated number of genes simultaneously. Tukey plots with mean shown by *. p-values from paired t-tests; **** = p <0.0001. Percentage of reads in three iPSC samples containing either K) indels, or L) translocations following delivery of editors targeting four genes simultaneously in combination with either a non-targeting (NTC) or p53 targeting siRNA. Analysed by targeted sequencing 7 days following delivery of editors. Data points represent variant frequencies at all on-target and known off-target sites of the four guide RNAs. Mean +/-SEM. Paired t-test analysis of differences between variant frequencies at individual targeted sites.

As iPSCs are continually undergoing DNA repair by error-free HDR (49) without adverse impact on their viability, we sought to better understand how the occurrence of DSBs during editing was related to p53 pathway activation by classifying nuclei according to the number of large and small foci. Whereas the presence of a single small γH2AX focus was sufficient to substantially increase the probability of high nuclear p53 levels compared to nuclei with no DNA damage foci in SpCas9 edited samples, this was not the case in either mock edited controls or nCas9-UGI:rAPO edited samples (Fig 3C, S4C). Rather, appreciable increases in the probability of high nuclear p53 levels were only observed in nuclei with 1 or more large, potentially more perdurant (63, 64), γH2AX foci in either mock edited controls or nCas9-UGI:rAPO edited samples (Fig 3D, S4D). Taken together we conclude from these data that despite an increase in the number of DSB foci associated with nCas9-UGI:rAPO editing compared to mock edited controls, the majority of these DNA repair events are of insufficient severity to trigger a p53 response. By contrast, DSBs generated by SpCas9 are markedly more likely to trigger a p53 DNA damage response.

To further understand whether DNA DSBs associated with nCas9-UGI:rAPO editing had a deleterious effect on the integrity of the genome we further characterised the nature of DNA DSBs by analysing 53BP1, which accumulates at γH2AX foci and promotes both error-prone NHEJ repair over HDR and chromosomal rearrangements by stabilising the double stranded ends of cleaved DNA strands (65, 66). 53BP1 levels within γH2AX foci were correlated with foci size (Fig 3E), indicating that large γH2AX foci generated by editing indeed represent a distinct class of DSBs undergoing error-prone, HDR-independent repair. Similar to large γH2AX foci, large 53BP1 foci were scarcely detectable in unedited controls, being primarily associated with SpCas9, and to a lesser extent with nCas9-UGI:rAPO multi-target edited samples (Fig 3F,G, S4E). Moreover, these large 53BP1 foci occurred more frequently in samples edited at multiple loci by SpCas9 than in single edited samples, further suggesting that these large foci were products of error-prone DSB-mediated editing (Fig 3G, S4E). As seen for γH2AX (Fig S4B), 53BP1 foci were also correlated with the quantity of non-targeting sgRNA delivered with nCas9-UGI:rAPO (Fig S4F), further confirming that DNA damage associated with nCas9-UGI:rAPO editing occurs independently of on-target editing.

Similar to observations for γH2AX, the accumulation of small 53BP1 foci had limited impact on the probability of high nuclear p53 levels in either mock edited controls or nCas9-UGI:rAPO edited samples (Fig 3H, S4G). By contrast, the presence of large foci increased the probability in both edited and unedited samples (Fig 3I, S4H), suggesting a qualitative difference between these two classes of DNA repair foci. The presence of large 53BP1 foci was most clearly correlated with elevated nuclear p53 levels in all edited and unedited sample groups (Fig 3C,D,H,I) consistent with the dual role of 53BP1 in both promoting error-prone repair and triggering p53 pathway activation (67). As large 53BP1 foci occurred most frequently in SpCas9 edited samples (Fig 3G, S4E) we therefore conclude that the differences in p53 activation observed between SpCas9 and base editing with nCas9-UGI:rAPO can be explained by the differences in the repair of DNA damage associated with each editing technology.

To begin to assess the severity of DNA damage associated with base editing we analysed the formation of micronuclei, which are known to be generated by Cas9 DSBs and can lead to complex structural rearrangements (chromothrypsis) when recombined with the genome (68, 69). Single target editing with nCas9-UGI:rAPO did not generate a detectable increase in micronuclei compared to those observed in mock edited controls, whereas a single SpCas9 edit was sufficient to increase the presence of detectable micronuclei (Fig 3J). Editing four targets simultaneously with nCas9-UGI:rAPO resulted in a small increase in micronuclei formation compared to either mock or single edited samples, whereas editing of four targets with SpCas9 resulted in a markedly greater increase (Fig 3J). The formation of micronuclei is therefore consistent with the observed differences in large DNA repair foci formation resulting from single or multiple edits with either platform, further suggesting that it is these DNA lesions that are likely to be genotoxic and therefore activate p53 mediated apoptosis.

p53 activity was shown to select against multiplexed gene knockout via NHEJ-mediated indel formation in SpCas9 edited samples (Fig 2C), raising the possibility that p53 mediated apoptosis may also serve to remove structural variants promoted by error-prone 53BP1-driven DNA repair. As the majority of DNA DSBs associated with nCas9-UGI:rAPO editing did not trigger p53 mediated cell cycle arrest or apoptosis this raised the possibility that these foci may also lead to indels or translocations, as has been suggested previously for other cytidine base editors (47), by evading detection by the p53 DNA damage sensing pathway.

To investigate this possibility, we employed targeted sequencing to analyse structural variants at the known on-and off-target loci targeted by the 4 sgRNAs. Consistent with our previous observations, we observed a significant increase in the occurrence of insertions and deletions at both sgRNA on-and off-target sites following transient p53 depletion in SpCa9 edited samples. By contrast, indels were undetectable in both control and p53 inhibited nCas9-UGI:rAPO edited samples (Fig 3K, S4I, Supplementary Table 7). Similarly, p53 inhibition increased the incidence of translocations between sgRNA target sites in SpCas9 edited samples whereas they remained undetectable in all nCas9-UGI:rAPO edited samples (Fig 3L, S4J, Supplementary Table 8).

Taking these data together we therefore conclude that whereas p53 activity acts to preserve the integrity of SpCas9 edited genomes by removing a proportion of undesirable chromosomal rearrangements alongside intended indels, it is not required to preserve the integrity of DNA repair following multiplexed editing with the nCas9-UGI:rAPO base editor.

### DNA damage response to nCas9-UGI:rAPO depends upon base editor module assembly

It has been demonstrated in other cellular contexts that both the nickase activity of the nCas9 component and insufficient activity of the UGI component of cytidine base editors can lead to DNA DSBs, p53 pathway activation, and cytotoxicity (44, 47). We therefore sought to understand if the low-level DNA damage and p53 pathway activation associated with multitarget base editing with nCas9-UGI:rAPO could be reduced to further improve the safety profile of the editor. We began by investigating how each component of the nCas9-UGI:rAPO editor contributes to the formation of DNA damage foci by assembling Pin-point cytidine base editors composed of distinct engineered SpCas9 modules (Fig 4A). Despite exhibiting the expected reductions in on-target editing efficiency and specificity (Fig S5A,B), neither the substitution of the nCas9-UGI component for a catalytically inactive dCas9-UGI, nor the omission of the UGI component resulted in the anticipated change (44, 47, 70) in either DSB formation (Fig 4B,C, S5C) or p53 pathway activation (Fig 4D,E, S5D). Rather, omission of the UGI component resulted in a reduction of the number of small 53BP1 and γH2AX foci to the basal levels observed in mock edited controls without affecting either large foci formation or p53 activity (Fig 4B-E), consistent with our previous observations that small DNA damage foci are not responsible for triggering the minimal p53 response associated with multiplexed nCas9-UGI:APO editing.

**Figure 4:**
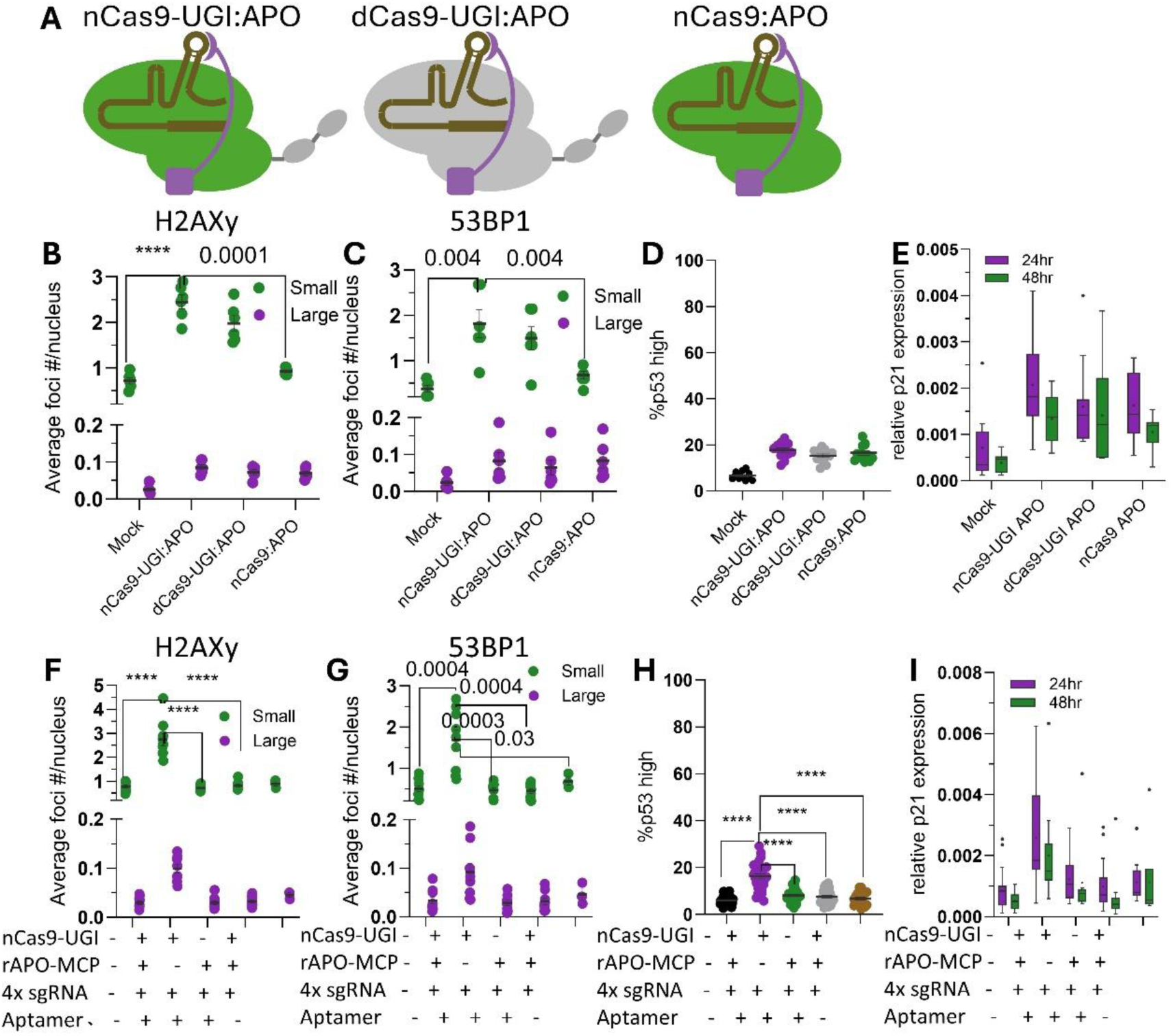
DNA damage response to nCas9-UGI:rAPO depends upon base editor module assembly. A) Schematic of the engineered SpCas9 variants. Quantification of the average number of B) γH2AX, or C) 53BP1 foci per nucleus classified as either large or small 24 hours following delivery of editors composed of rAPO and the indicated engineered SpCas9 variant targeting four genes simultaneously. Mean +/-SEM. p-values from paired t-tests; **** = p<0.0001. Quantification of D) the percentage of nuclei classified as containing high levels of p53 by microscopy (Mean +/-SEM), or E) p21 expression in mock edited samples, or samples edited with an editor composed of rAPO and the indicated engineered SpCas9 variant (Tukey plots with mean shown by *). Analysed D) 24 hours, or E) at the indicated time following delivery of the editor. Quantification of the average number of F) γH2AX, or G) 53BP1 foci per nucleus classified as either large or small 24 hours following delivery of the indicated components of the nCas9-UGI:rAPO editor in combination with sgRNAs targeting four genes simultaneously. MS2-less indicates omission of the MS2 aptamer required for assembly of the nCas9-UGI:rAPO editor from all four sgRNAs. Mean +/-SEM. p-values from paired t-tests; **** = p<0.0001. Quantification of H) the percentage of nuclei classified as containing high levels of p53 by microscopy (mean +/-SEM, p-values from paired t-tests; **** = p<0.0001), or I) p21 expression following delivery of the indicated components of the nCas9-UGI:rAPO editor in combination with sgRNAs targeting four genes simultaneously (Tukey plots with mean shown by *). Analysed H) 24hrs, or I) at the indicated time following delivery of the editor. MS2-less indicates omission of the MS2 aptamer required for assembly of the nCas9-UGI:rAPO editor from all four sgRNAs.

As UNG inhibition inhibits excision of deaminated cytidine bases this suggested that the low-severity DNA damage attributed to the UGI component may be due to its interaction with uracil bases generated by the APOBEC1 component. In support of this hypothesis, when either the nCas9-UGI or APOBEC1 components were introduced separately in combination with the four Pin-point sgRNAs, both γH2AX and 53BP1 foci (Fig 4F,G) and p53 activity were returned to basal levels (Fig 4H,I). Moreover, abolishing assembly of the base editor complex by omitting the aptamer component from the four sgRNAs also restored DNA repair and p53 activity to basal levels (Fig 4F-I). Taken together these data demonstrate that DNA DSB formation by nCas9-UGI:rAPO depends on the localised activity of both the deaminase and UGI modules of the base editor.

### Optimised module assembly eliminates DNA damage response to nCas9-UGI:rAPO multiplexed editing

Neither nCas9-UGI nor untethered APOBEC1 alone cause detectable DNA damage or p53 pathway activity (Fig 4F-I). Moreover, the nCas9-UGI:rAPO complex only resulted in detectable DNA damage when delivered with larger sgRNA quantities (Fig 3B,G, S4B, F). We therefore reasoned that it may be possible to further improve the balance of DNA DSB formation to efficient on-target editing by modulating the assembly of the base editor via limiting the availability of targeting sgRNA. As observed previously in comparisons between single and multiplexed editing, DNA damage foci formation and p53 activity showed a dose dependent response to sgRNA quantity, returning to levels comparable to mock edited controls when all 4 targeting sgRNAs were reduced from 40pM to 10pM (160pM total sgRNA to 40pM) (Fig 5A-C). Reduction of individual sgRNA quantity from 40pM to 10pM minimally impacted editing by the two most efficient sgRNAs, whereas the less active sgRNAs showed a greater sensitivity to sgRNA quantity (Fig 5D). Thus, given sufficiently optimised sgRNA designs the low-level DNA damage associated with multiplexed editing with the nCas9-UGI:rAPO editor can be eliminated without significantly reducing editing efficiency. Taken together these data highlight that the modularity of the Pin-point platform allows cytidine base editor activity to be tuned via the quantities of each deaminase recruiting guide RNA component, creating opportunities to optimise the combined safety and efficacy profile of base editors formulated for specific single target or multi target editing applications.

**Figure 5:**
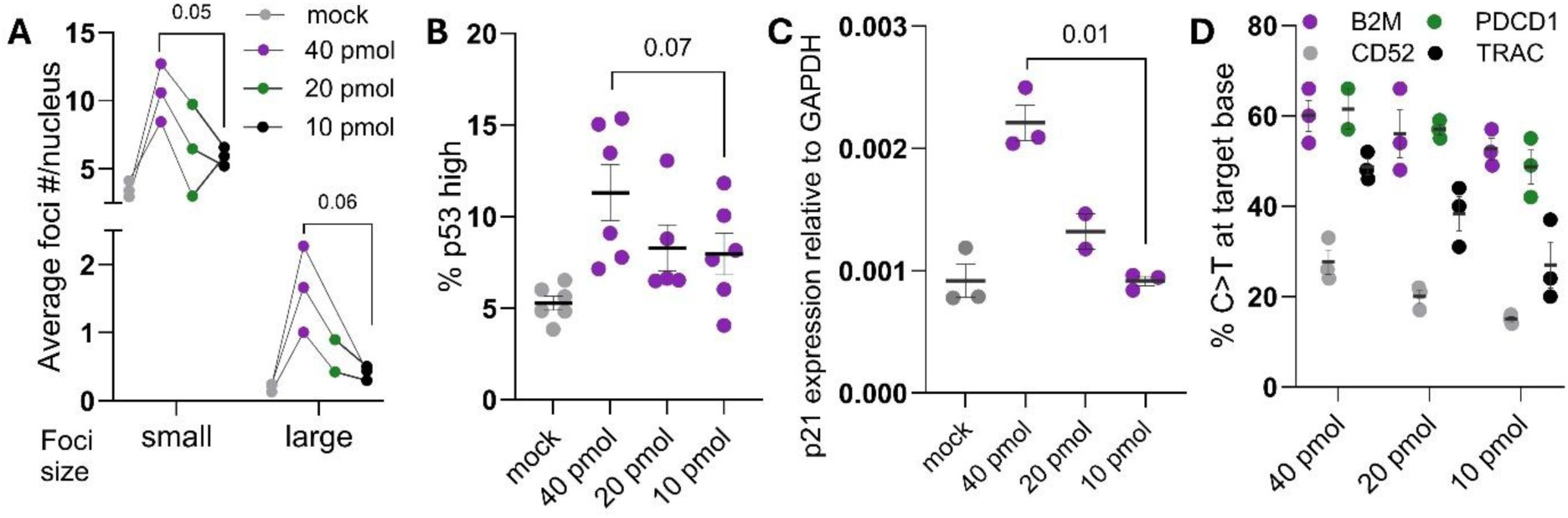
Optimised module assembly minimises DNA damage response to nCas9-UGI:rAPO multiplexed editing. Quantification of the average number of A) γH2AX foci per nucleus classified as either large or small 24 hours following delivery of the nCas9-UGI:rAPO editor in combination with the indicated quantities of each of the sgRNAs targeting four genes simultaneously. p-values from paired t-tests. Connecting lines show paired measurements from n=3 iPSC lines. Quantification of B) the percentage of nuclei classified as containing high levels of p53 by microscopy, or C) p21 expression following delivery of the nCas9-UGI:rAPO editor in combination with the indicated quantities of each of the sgRNAs targeting four genes simultaneously, analysed 24 hours following delivery of the editor. Mean +/-SEM, p-values from paired t-tests. D) Percentage of base conversion at the target nucleotide or four genes edited simultaneously following delivery of the nCas9-UGI:rAPO editor in combination with the indicated quantities of each of the sgRNAs. Editing analysed by Sanger sequencing 7 days following delivery of editor formulations. Mean +/-SEM.

## DISCUSSION

Here we present our finding that efficient base editing can be achieved in DNA damage sensitive iPSCs by transient expression of a modular cytidine base editor via delivery of an optimised RNA formulation. In both single target and multiplexed gene knockout applications the cytidine base editor induced markedly reduced levels of p53 pathway activity compared to editing with SpCas9 nuclease, resulting in increased viability of edited cells and improved multiplexed editing efficiency. Whereas both p53 pathway activity and associated reduction in viable edited cell yields was correlated with the number of DNA edits at either on-target or partially homologous off-target sites by Cas9 nuclease, this was not the case for base editing. Rather, the low levels of p53 pathway induction observed when base editing multiple sites was independent of the number of simultaneous on-target editing events. These favourable attributes of Cas9-based base editors create opportunities for the efficient manufacturing of genetically enhanced allogeneic iPSC-derived cell therapies through multiplexed gene knockout in combination with rational engineering of cellular phenotypes (71–73). Moreover, the ability of base editing to precisely modify disease-causing genetic variants with minimal adverse impacts on stem cell fitness may support future developments in autologous stem cell therapies (74, 75).

In addition to enhancing editing efficiency, the reduced p53-activating DNA damage associated with multiplexed base editing also presents a number of safety advantages compared to nuclease editing when edited iPSC products are intended for therapeutic application. Whereas multiplexed nuclease editing resulted in anticipated genome rearrangements in addition to indels at both on-target and previously identified off-target sites, these were undetectable following multiplexed base editing in contrast to a previous reports in hematopoietic stem cells (47). In line with the role of p53 as the guardian of genome integrity we show that p53 mediated apoptosis selects against both intended on-target nuclease editing outcomes in addition to unintended off-target editing and genome rearrangement. However, in contrast to previous reports we did not observe enhanced base editing efficiency or any increase in undesirable editing outcomes when p53 activity was reduced (32, 33, 47). Importantly, as p53 activity selects against on-target nuclease editing this raises the question of whether there are heritable, potentially oncogenic, differences between edited clones that escape selection and those that are eliminated by apoptosis.

While it is possible to sequence edited master cell banks to identify variants within known DNA damage sensing and tumour suppressor genes, nuclease editing may nonetheless select for variants either in genes unknown to influence the response to DNA damage, or in incompletely understood regulatory regions of the genome. Moreover, it remains unclear to what extent the greater p53 selection associated with multiplexed nuclease editing is ameliorated by multiple rounds of single target editing. We therefore propose that in addition to reducing known risks to genome stability, base editing may also reduce the risk of selecting for genetic variants related to less well understood processes governing the survival of edited clones.

While it is acknowledged that our fixed sample microscopy analysis of editing induced DNA damage and repair requires cautious interpretation, we nonetheless propose that the differences in foci size and frequency observed at a snapshot in time reflect underlying differences in DNA repair foci dynamics. The initial, rapid accumulation of phosphorylated H2AX flanking DNA double strand breaks results in the formation of large γH2AX foci, which subsequently dissipate over an extended period of time following repair (63, 64). Thus, the relative frequencies of large foci induced by editing serves as a comparative metric of ongoing, unresolved DNA repair, whereas the frequency of small foci serves primarily as a metric of previously completed repair events. Consistent with this interpretation, high levels of p53 tended to accumulate in nuclei with large foci, indicating ongoing DNA damage signalling. Moreover, large SpCas9-induced γH2AX foci tended to accumulate high levels of the NHEJ promoting factor 53BP1, indicating that these unresolved DNA damage foci undergo error-prone repair. By contrast, foci with large accumulations of 53BP1 were barely detectable in base-edited samples, consistent with the undetectable levels of indels or interchromosomal translocations. Small 53BP1 foci were nonetheless detectable at greater frequencies in base edited samples than in nuclease edited samples. However, these large numbers of foci containing small amounts of 53BP1 were insufficient to trigger a p53 DNA damage response or generate detectable levels of imprecise DNA repair, consistent with these processes depending on the relative amount of NHEJ and HDR promoting factors *at individual repair foci* (67) rather than their absolute levels in the nucleus. We therefore highlight the importance of basing interpretation on quantitative analysis of repair factors within qualitatively different classes of foci rather than bulk measurement. To our knowledge the distinct DNA damage profiles of nuclease and base editors have not been resolved at this detail to date, and we therefore expect the results we present here to inform design and interpretation of future studies into DNA damage and repair associated with genome editing.

Single nucleus imaging of the DNA damage response to DNA editing with nuclease and base editors therefore clarifies the seemingly paradoxical previous reports that Cas9 nickase-based cytidine base editors induce comparable levels of γH2AX labelled DNA DSBs as Cas9 nuclease, despite generating markedly fewer permanent insertions or deletions in the genome (44, 46). Our foci-level analysis demonstrates both qualitative and quantitative differences between the two editing modalities; whereas SpCas9 nuclease primarily induces p53 pathway-activating large γH2AX labelled DNA repair foci, the modular cytidine base editor composed of nCas-UGI and Rat APOBEC1 predominantly induces formation of multiple small foci, consistent with the similar aggregate γH2AX levels observed for each editor type derived from either per-cell or bulk measurements (44, 46). As pluripotent cells are continually undergoing and repairing DNA damage due to normal replicative stress (49) without triggering a p53 response, we speculate that base editor-induced DNA repair foci more closely resemble these benign lesions than Cas9 induced DSBs.

By combining the modularity of our base editor platform with single nucleus measurements of the DNA damage response we were able to determine that the introduction of single strand breaks had no detectable effect on the frequency of p53 pathway-activating DNA DSB repair foci, suggesting that cytidine base editors incorporating Cas nickase modules are not intrinsically more genotoxic in contrast to conclusions of previous reports (44, 47). Similarly, the small, likely short-lived, DNA repair foci attributed to the activity of the UGI component also did not activate a p53 response, suggesting that this class of DNA damage events is repaired within the normal course of genome maintenance occurring in cycling pluripotent cells (49). Rather, the class of p53 activating DNA damage events we detected at low frequency with our modular CBE occurred as a result of recruiting Rat APOBEC1 to a Cas9 bound genomic locus, irrespective of whether a single strand DNA break was introduced or BER was inhibited by the UGI component. This finding, that transient expression of the APOBEC1 module by mRNA delivery is not intrinsically genotoxic, is in contrast to previous reports of the activity of other APOBEC family members (76) and further highlights the potential for safe application of the modular base editing platform. It has been reported previously that insufficient inhibition of UNG activity by UGI results in p53 mediated genotoxicity in hematopoietic stem cells (47), which are also highly sensitive to DNA damage, however it remains unclear why we did not detect a similar requirement for UGI in pluripotent stem cells. We speculate that the discrepancies we observed with previous studies may be explained either by differences in the quantities of editor delivered by mRNA, or the duration of expression of editors and sgRNAs delivered by plasmid compared to RNA electroporation as used in this study.

By dissecting the activities of each component of the modular CBE using single nucleus and single foci measurements we determined that p53 activating DNA damage events depended upon the assembly of the base editing complex via the sgRNA-encoded aptamer, further demonstrating that transient expression of optimised quantities of either the nCas-UGI or APOBEC1 deaminase components alone is not genotoxic. This discovery uncovered the opportunity to further optimise the safety profile of modular base editors by fine tuning base editor complex assembly via the aptamer. As proof of concept of the potential of tuning base editor assembly we demonstrate that p53 activating DNA damage can be minimised by limiting the quantity of aptamer encoding sgRNA delivered without materially impacting editing efficiency, raising the possibility of developing this strategy to further improve the safety profile of base editors. Taken together with the ever-expanding repertoire of novel deaminases created by rational engineering, directed evolution, and generative AI (77–81) we envision that the modular aptamer recruitment platform will enhance the potential to further imporove the safety and efficacy profile of base editors for advanced cell therapy manufacturing.

## DATA AVAILABILITY

The NGS datasets reported in this manuscript are available on Sequence Read Archive with BioProject ID PRJNA1272047 https://www.ncbi.nlm.nih.gov/bioproject/PRJNA1272047

Supplementary Data are available online

## AUTHOR CONTRIBUTIONS

Robert Blassberg: Conceptualization, Supervision, Investigation, Methodology, Formal analysis, Visualisation, Writing—original draft.

Olga Mielczarek: Investigation, Methodology, Formal analysis

Jesse Stombaugh: Data curation, Software, Formal analysis

Hanneke Okkenhaug: Investigation, Data curation, Formal analysis

Alice Abreu Torres: Investigation

Paul Russell: Conceptualisation, Investigation

John Lambourne: Project administration

Immacolata Porreca: Conceptualization, Project administration, Writing—review & editing

## ACKNOWLEDGEMENTS

We thank Marcela Buricova, Alexis Duringer, Leigh-Anne Thomas and Amanda Haupt for comments on the manuscript and for their helpful discussion and support.

## REFERENCES

1. Kirkeby A., Main H. and Carpenter M. (2025) Pluripotent stem-cell-derived therapies in clinical trial: A 2025 update. Cell Stem Cell, 32, 10–37.

2. Blassberg R. (2022) Genome Editing of Pluripotent Stem Cells for Adoptive and Regenerative Cell Therapies. GEN Biotechnology, 1, 77–90.

3. Wood K.J. and Goto R. (2012) Mechanisms of Rejection: Current Perspectives. Transplantation, 93, 1.

4. Li Q. and Lan P. (2023) Activation of immune signals during organ transplantation. Sig Transduct Target Ther, 8, 1–26.

5. Felix N.J. and Allen P.M. (2007) Specificity of T-cell alloreactivity. Nat Rev Immunol, 7, 942–953.

6. Gumber D. and Wang L.D. (2022) Improving CAR-T immunotherapy: Overcoming the challenges of T cell exhaustion. eBioMedicine, 77.

7. Hu X., Manner K., DeJesus R., White K., Gattis C., Ngo P., Bandoro C., Tham E., Chu E.Y., Young C., et al. (2023) Hypoimmune anti-CD19 chimeric antigen receptor T cells provide lasting tumor control in fully immunocompetent allogeneic humanized mice. Nat Commun, 14, 2020.

8. Kagoya Y., Guo T., Yeung B., Saso K., Anczurowski M., Wang C.-H., Murata K., Sugata K., Saijo H., Matsunaga Y., et al. (2020) Genetic Ablation of HLA Class I, Class II, and the T-cell Receptor Enables Allogeneic T Cells to Be Used for Adoptive T-cell Therapy. Cancer Immunol Res, 8, 926–936.

9. Benjamin R., Graham C., Yallop D., Jozwik A., Mirci-Danicar O.C., Lucchini G., Pinner D., Jain N., Kantarjian H., Boissel N., et al. (2020) Genome-edited, donor-derived allogeneic anti-CD19 chimeric antigen receptor T cells in paediatric and adult B-cell acute lymphoblastic leukaemia: results of two phase 1 studies. The Lancet, 396, 1885–1894.

10. Ottaviano G., Georgiadis C., Gkazi S.A., Syed F., Zhan H., Etuk A., Preece R., Chu J., Kubat A., Adams S., et al. (2022) Phase 1 clinical trial of CRISPR-engineered CAR19 universal T cells for treatment of children with refractory B cell leukemia. Science Translational Medicine, 14, eabq3010.

11. Stadtmauer E.A., Fraietta J.A., Davis M.M., Cohen A.D., Weber K.L., Lancaster E., Mangan P.A., Kulikovskaya I., Gupta M., Chen F., et al. (2020) CRISPR-engineered T cells in patients with refractory cancer. Science, 367, eaba7365.

12. Frangoul H., Altshuler D., Cappellini M.D., Chen Y.-S., Domm J., Eustace B.K., Foell J., Fuente J. de la, Grupp S., Handgretinger R., et al. (2021) CRISPR-Cas9 Gene Editing for Sickle Cell Disease and β-Thalassemia. New England Journal of Medicine, 384, 252–260.

13. Adikusuma F., Piltz S., Corbett M.A., Turvey M., McColl S.R., Helbig K.J., Beard M.R., Hughes J., Pomerantz R.T. and Thomas P.Q. (2018) Large deletions induced by Cas9 cleavage. Nature, 560, E8–E9.

14. Alanis-Lobato G., Zohren J., McCarthy A., Fogarty N.M.E., Kubikova N., Hardman E., Greco M., Wells D., Turner J.M.A. and Niakan K.K. (2021) Frequent loss of heterozygosity in CRISPR-Cas9–edited early human embryos. PNAS, 118.

15. Kosicki M., Tomberg K. and Bradley A. (2018) Repair of double-strand breaks induced by CRISPR– Cas9 leads to large deletions and complex rearrangements. Nat Biotechnol, 36, 765–771.

16. Cullot G., Boutin J., Toutain J., Prat F., Pennamen P., Rooryck C., Teichmann M., Rousseau E., Lamrissi-Garcia I., Guyonnet-Duperat V., et al. (2019) CRISPR-Cas9 genome editing induces megabase-scale chromosomal truncations. Nat Commun, 10, 1136.

17. Nahmad A.D., Reuveni E., Goldschmidt E., Tenne T., Liberman M., Horovitz-Fried M., Khosravi R., Kobo H., Reinstein E., Madi A., et al. (2022) Frequent aneuploidy in primary human T cells after CRISPR–Cas9 cleavage. Nat Biotechnol, 40, 1807–1813.

18. Ousterout D.G., Kabadi A.M., Thakore P.I., Majoros W.H., Reddy T.E. and Gersbach C.A. (2015) Multiplex CRISPR/Cas9-based genome editing for correction of dystrophin mutations that cause Duchenne muscular dystrophy. Nat Commun, 6, 6244.

19. Samuelson C., Radtke S., Zhu H., Llewellyn M., Fields E., Cook S., Huang M.-L.W., Jerome K.R., Kiem H.-P. and Humbert O. (2021) Multiplex CRISPR/Cas9 genome editing in hematopoietic stem cells for fetal hemoglobin reinduction generates chromosomal translocations. Mol Ther Methods Clin Dev, 23, 507–523.

20. Webber B.R., Lonetree C., Kluesner M.G., Johnson M.J., Pomeroy E.J., Diers M.D., Lahr W.S., Draper G.M., Slipek N.J., Smeester B.A., et al. (2019) Highly efficient multiplex human T cell engineering without double-strand breaks using Cas9 base editors. Nat Commun, 10, 5222.

21. Williams A.B. and Schumacher B. (2016) p53 in the DNA-Damage-Repair Process. Cold Spring Harb Perspect Med, 6, a026070.

22. Dumitru R., Gama V., Fagan B.M., Bower J.J., Swahari V., Pevny L.H. and Deshmukh M. (2012) Human Embryonic Stem Cells Have Constitutively Active Bax at the Golgi and Are Primed to Undergo Rapid Apoptosis. Molecular Cell, 46, 573–583.

23. Heyer B.S., MacAuley A., Behrendtsen O. and Werb Z. (2000) Hypersensitivity to DNA damage leads to increased apoptosis during early mouse development. Genes Dev., 14, 2072–2084.

24. Ihry R.J., Worringer K.A., Salick M.R., Frias E., Ho D., Theriault K., Kommineni S., Chen J., Sondey M., Ye C., et al. (2018) p53 inhibits CRISPR–Cas9 engineering in human pluripotent stem cells. Nat Med, 24, 939–946.

25. Merkle F.T., Ghosh S., Kamitaki N., Mitchell J., Avior Y., Mello C., Kashin S., Mekhoubad S., Ilic D., Charlton M., et al. (2017) Human pluripotent stem cells recurrently acquire and expand dominant negative P53 mutations. Nature, 545, 229–233.

26. Enache O.M., Rendo V., Abdusamad M., Lam D., Davison D., Pal S., Currimjee N., Hess J., Pantel S., Nag A., et al. (2020) Cas9 activates the p53 pathway and selects for p53-inactivating mutations. Nat Genet, 52, 662–668.

27. Jiang L., Ingelshed K., Shen Y., Boddul S.V., Iyer V.S., Kasza Z., Sedimbi S., Lane D.P. and Wermeling F. (2022) CRISPR/Cas9-Induced DNA Damage Enriches for Mutations in a p53-Linked Interactome: Implications for CRISPR-Based Therapies. Cancer Research, 82, 36–45.

28. Sinha S., Barbosa K., Cheng K., Leiserson M.D.M., Jain P., Deshpande A., Wilson D.M., Ryan B.M., Luo J., Ronai Z.A., et al. (2021) A systematic genome-wide mapping of oncogenic mutation selection during CRISPR-Cas9 genome editing. Nat Commun, 12, 6512.

29. Bai Q., Ramirez J.-M., Becker F., Pantesco V., Lavabre-Bertrand T., Hovatta O., Lemaître J.-M., Pellestor F. and De Vos J. (2015) Temporal Analysis of Genome Alterations Induced by Single-Cell Passaging in Human Embryonic Stem Cells. Stem Cells Dev, 24, 653–662.

30. Assou S., Girault N., Plinet M., Bouckenheimer J., Sansac C., Combe M., Mianné J., Bourguignon C., Fieldes M., Ahmed E., et al. (2020) Recurrent Genetic Abnormalities in Human Pluripotent Stem Cells: Definition and Routine Detection in Culture Supernatant by Targeted Droplet Digital PCR. Stem Cell Reports, 14, 1–8.

31. Sürün D., Schneider A., Mircetic J., Neumann K., Lansing F., Paszkowski-Rogacz M., Hänchen V., Lee-Kirsch M.A. and Buchholz F. (2020) Efficient Generation and Correction of Mutations in Human iPS Cells Utilizing mRNAs of CRISPR Base Editors and Prime Editors. Genes, 11, 511.

32. Park J.-C., Kim Y.-J., Hwang G.-H., Kang C.Y., Bae S. and Cha H.-J. (2024) Enhancing genome editing in hPSCs through dual inhibition of DNA damage response and repair pathways. Nat Commun, 15, 4002.

33. Li M., Zhong A., Wu Y., Sidharta M., Beaury M., Zhao X., Studer L. and Zhou T. (2022) Transient inhibition of p53 enhances prime editing and cytosine base-editing efficiencies in human pluripotent stem cells. Nat Commun, 13, 6354.

34. Park J.-C., Jang H.-K., Kim J., Han J.H., Jung Y., Kim K., Bae S. and Cha H.-J. (2022) High expression of uracil DNA glycosylase determines C to T substitution in human pluripotent stem cells. Molecular Therapy - Nucleic Acids, 27, 175–183.

35. Li H., Busquets O., Verma Y., Syed K.M., Kutnowski N., Pangilinan G.R., Gilbert L.A., Bateup H.S., Rio D.C., Hockemeyer D., et al. (2022) Highly efficient generation of isogenic pluripotent stem cell models using prime editing. eLife, 11, e79208.

36. Anzalone A.V., Koblan L.W. and Liu D.R. (2020) Genome editing with CRISPR–Cas nucleases, base editors, transposases and prime editors. Nat Biotechnol, 38, 824–844.

37. Rees H.A. and Liu D.R. (2018) Base editing: precision chemistry on the genome and transcriptome of living cells. Nat Rev Genet, 19, 770–788.

38. Komor A.C., Kim Y.B., Packer M.S., Zuris J.A. and Liu D.R. (2016) Programmable editing of a target base in genomic DNA without double-stranded DNA cleavage. Nature, 533, 420–424.

39. Gaudelli N.M., Lam D.K., Rees H.A., Solá-Esteves N.M., Barrera L.A., Born D.A., Edwards A., Gehrke J.M., Lee S.-J., Liquori A.J., et al. (2020) Directed evolution of adenine base editors with increased activity and therapeutic application. Nat Biotechnol, 38, 892–900.

40. Kluesner M.G., Lahr W.S., Lonetree C., Smeester B.A., Qiu X., Slipek N.J., Claudio Vázquez P.N., Pitzen S.P., Pomeroy E.J., Vignes M.J., et al. (2021) CRISPR-Cas9 cytidine and adenosine base editing of splice-sites mediates highly-efficient disruption of proteins in primary and immortalized cells. Nat Commun, 12, 2437.

41. Billon P., Bryant E.E., Joseph S.A., Nambiar T.S., Hayward S.B., Rothstein R. and Ciccia A. (2017) CRISPR-Mediated Base Editing Enables Efficient Disruption of Eukaryotic Genes through Induction of STOP Codons. Molecular Cell, 67, 1068–1079.e4.

42. Kuscu C., Parlak M., Tufan T., Yang J., Szlachta K., Wei X., Mammadov R. and Adli M. (2017) CRISPR-STOP: gene silencing through base-editing-induced nonsense mutations. Nat Methods, 14, 710–712.

43. Komor A.C., Zhao K.T., Packer M.S., Gaudelli N.M., Waterbury A.L., Koblan L.W., Kim Y.B., Badran A.H. and Liu D.R. (2017) Improved base excision repair inhibition and bacteriophage Mu Gam protein yields C:G-to-T:A base editors with higher efficiency and product purity. Science Advances, 3, eaao4774.

44. Wang X., Ding C., Yu W., Wang Y., He S., Yang B., Xiong Y.-C., Wei J., Li J., Liang J., et al. (2020) Cas12a Base Editors Induce Efficient and Specific Editing with Low DNA Damage Response. Cell Reports, 31.

45. Zhang S., Yuan B., Cao J., Song L., Chen J., Qiu J., Qiu Z., Zhao X.-M., Chen J. and Cheng T.-L. (2023) TadA orthologs enable both cytosine and adenine editing of base editors. Nat Commun, 14, 414.

46. Yuan B., Zhang S., Song L., Chen J., Cao J., Qiu J., Qiu Z., Chen J., Zhao X.-M. and Cheng T.-L. (2023) Engineering of cytosine base editors with DNA damage minimization and editing scope diversification. Nucleic Acids Research, 51, e105.

47. Fiumara M., Ferrari S., Omer-Javed A., Beretta S., Albano L., Canarutto D., Varesi A., Gaddoni C., Brombin C., Cugnata F., et al. (2023) Genotoxic effects of base and prime editing in human hematopoietic stem cells. Nat Biotechnol, 10.1038/s41587-023-01915-4.

48. Ahuja A.K., Jodkowska K., Teloni F., Bizard A.H., Zellweger R., Herrador R., Ortega S., Hickson I.D., Altmeyer M., Mendez J., et al. (2016) A short G1 phase imposes constitutive replication stress and fork remodelling in mouse embryonic stem cells. Nat Commun, 7, 10660.

49. Vallabhaneni H., Lynch P.J., Chen G., Park K., Liu Y., Goehe R., Mallon B.S., Boehm M. and Hursh D.A. (2018) High Basal Levels of γH2AX in Human Induced Pluripotent Stem Cells Are Linked to Replication-Associated DNA Damage and Repair. STEM CELLS, 36, 1501–1513.

50. Desmarais J.A., Hoffmann M.J., Bingham G., Gagou M.E., Meuth M. and Andrews P.W. (2012) Human Embryonic Stem Cells Fail to Activate CHK1 and Commit to Apoptosis in Response to DNA Replication Stress. STEM CELLS, 30, 1385–1393.

51. Collantes J.C., Tan V.M., Xu H., Ruiz-Urigüen M., Alasadi A., Guo J., Tao H., Su C., Tyc K.M., Selmi T., et al. (2021) Development and Characterization of a Modular CRISPR and RNA Aptamer Mediated Base Editing System. The CRISPR Journal, 4, 58–68.

52. Porreca I., Blassberg R., Harbottle J., Joubert B., Mielczarek O., Stombaugh J., Hemphill K., Sumner J., Pazeraitis D., Touza J.L., et al. (2024) An aptamer-mediated base editing platform for simultaneous knockin and multiple gene knockout for allogeneic CAR-T cells generation. Molecular Therapy, 32, 2692–2710.

53. Burridge P.W., Thompson S., Millrod M.A., Weinberg S., Yuan X., Peters A., Mahairaki V., Koliatsos V.E., Tung L. and Zambidis E.T. (2011) A Universal System for Highly Efficient Cardiac Differentiation of Human Induced Pluripotent Stem Cells That Eliminates Interline Variability. PLOS ONE, 6, e18293.

54. Kreitzer F.R., Salomonis N., Sheehan A., Huang M., Park J.S., Spindler M.J., Lizarraga P., Weiss W.A., So P.-L. and Conklin B.R. (2013) A robust method to derive functional neural crest cells from human pluripotent stem cells. Am J Stem Cells, 2, 119–131.

55. Jinek M., Chylinski K., Fonfara I., Hauer M., Doudna J.A. and Charpentier E. (2012) A Programmable Dual-RNA–Guided DNA Endonuclease in Adaptive Bacterial Immunity. Science, 337, 816–821.

56. Xu L., Liu Y. and Han R. (2019) BEAT: A Python Program to Quantify Base Editing from Sanger Sequencing. The CRISPR Journal, 2, 223–229.

57. Brinkman E.K., Chen T., Amendola M. and van Steensel B. (2014) Easy quantitative assessment of genome editing by sequence trace decomposition. Nucleic Acids Research, 42, e168.

58. Miller N.A., Farrow E.G., Gibson M., Willig L.K., Twist G., Yoo B., Marrs T., Corder S., Krivohlavek L., Walter A., et al. (2015) A 26-hour system of highly sensitive whole genome sequencing for emergency management of genetic diseases. Genome Medicine, 7, 1–16.

59. Chen X., Schulz-Trieglaff O., Shaw R., Barnes B., Schlesinger F., Källberg M., Cox A.J., Kruglyak S. and Saunders C.T. (2016) Manta: Rapid detection of structural variants and indels for germline and cancer sequencing applications. Bioinformatics, 32, 1220–1222.

60. van den Berg J., G. Manjón A., Kielbassa K., Feringa F.M., Freire R. and Medema R.H. (2018) A limited number of double-strand DNA breaks is sufficient to delay cell cycle progression. Nucleic Acids Research, 46, 10132–10144.

61. Singh A., Smedley G.D., Rose J.-G., Fredriksen K., Zhang Y., Li L. and Yuan S.H. (2024) A high efficiency precision genome editing method with CRISPR in iPSCs. Sci Rep, 14, 9933.

62. Mah L.-J., El-Osta A. and Karagiannis T.C. (2010) γH2AX: a sensitive molecular marker of DNA damage and repair. Leukemia, 24, 679–686.

63. Rogakou E.P., Pilch D.R., Orr A.H., Ivanova V.S. and Bonner W.M. (1998) DNA Double-stranded Breaks Induce Histone H2AX Phosphorylation on Serine 139 *. Journal of Biological Chemistry, 273, 5858–5868.

64. Rothkamm K., Barnard S., Moquet J., Ellender M., Rana Z. and Burdak-Rothkamm S. (2015) DNA damage foci: Meaning and significance. Environmental and Molecular Mutagenesis, 56, 491– 504.

65. Mirman Z. and Lange T. de (2020) 53BP1: a DSB escort. Genes Dev., 34, 7–23.

66. Rappold I., Iwabuchi K., Date T. and Chen J. (2001) Tumor Suppressor P53 Binding Protein 1 (53bp1) Is Involved in DNA Damage–Signaling Pathways. Journal of Cell Biology, 153, 613– 620.

67. Cuella-Martin R., Oliveira C., Lockstone H.E., Snellenberg S., Grolmusova N. and Chapman J.R. (2016) 53BP1 Integrates DNA Repair and p53-Dependent Cell Fate Decisions via Distinct Mechanisms. Molecular Cell, 64, 51–64.

68. Leibowitz M.L., Papathanasiou S., Doerfler P.A., Blaine L.J., Sun L., Yao Y., Zhang C.-Z., Weiss M.J. and Pellman D. (2021) Chromothripsis as an on-target consequence of CRISPR–Cas9 genome editing. Nat Genet, 53, 895–905.

69. Zhang C.-Z., Spektor A., Cornils H., Francis J.M., Jackson E.K., Liu S., Meyerson M. and Pellman D. (2015) Chromothripsis from DNA damage in micronuclei. Nature, 522, 179–184.

70. Bregenhorn S., Kallenberger L., Artola-Borán M., Peña-Diaz J. and Jiricny J. (2016) Non-canonical uracil processing in DNA gives rise to double-strand breaks and deletions: relevance to class switch recombination. Nucleic Acids Research, 44, 2691–2705.

71. Zeng J., Wu Y., Ren C., Bonanno J., Shen A.H., Shea D., Gehrke J.M., Clement K., Luk K., Yao Q., et al. (2020) Therapeutic base editing of human hematopoietic stem cells. Nat Med, 26, 535– 541.

72. Casirati G., Cosentino A., Mucci A., Salah Mahmoud M., Ugarte Zabala I., Zeng J., Ficarro S.B., Klatt D., Brendel C., Rambaldi A., et al. (2023) Epitope editing enables targeted immunotherapy of acute myeloid leukaemia. Nature, 621, 404–414.

73. Wang J., Wang K., Deng Z., Zhong Z., Sun G., Mei Q., Zhou F., Deng Z. and Sun Y. (2024) Engineered cytosine base editor enabling broad-scope and high-fidelity gene editing in Streptomyces. Nat Commun, 15, 5687.

74. Bzhilyanskaya V., Ma L., Liu S., Fox L.R., Whittaker M.N., Meis R.J., Choi U., Lawson A., Ma M., Theobald N., et al. (2024) High-fidelity PAMless base editing of hematopoietic stem cells to treat chronic granulomatous disease. Science Translational Medicine, 16, eadj6779.

75. Newby G.A., Yen J.S., Woodard K.J., Mayuranathan T., Lazzarotto C.R., Li Y., Sheppard-Tillman H., Porter S.N., Yao Y., Mayberry K., et al. (2021) Base editing of haematopoietic stem cells rescues sickle cell disease in mice. Nature, 595, 295–302.

76. Landry S., Narvaiza I., Linfesty D.C. and Weitzman M.D. (2011) APOBEC3A can activate the DNA damage response and cause cell-cycle arrest. EMBO reports, 12, 444–450.

77. Yang L., Huo Y., Wang M., Zhang D., Zhang T., Wu H., Rao X., Meng H., Yin S., Mei J., et al. (2024) Engineering APOBEC3A deaminase for highly accurate and efficient base editing. Nat Chem Biol, 20, 1176–1187.

78. Lee S., Ding N., Sun Y., Yuan T., Li J., Yuan Q., Liu L., Yang J., Wang Q., Kolomeisky A.B., et al. (2020) Single C-to-T substitution using engineered APOBEC3G-nCas9 base editors with minimum genome-and transcriptome-wide off-target effects. Science Advances, 6, eaba1773.

79. Kim Y.B., Komor A.C., Levy J.M., Packer M.S., Zhao K.T. and Liu D.R. (2017) Increasing the genome-targeting scope and precision of base editing with engineered Cas9-cytidine deaminase fusions. Nat Biotechnol, 35, 371–376.

80. Richter M.F., Zhao K.T., Eton E., Lapinaite A., Newby G.A., Thuronyi B.W., Wilson C., Koblan L.W., Zeng J., Bauer D.E., et al. (2020) Phage-assisted evolution of an adenine base editor with improved Cas domain compatibility and activity. Nat Biotechnol, 38, 883–891.

81. Ruffolo J.A., Nayfach S., Gallagher J., Bhatnagar A., Beazer J., Hussain R., Russ J., Yip J., Hill E., Pacesa M., et al. (2024) Design of highly functional genome editors by modeling the universe of CRISPR-Cas sequences. 10.1101/2024.04.22.590591.

