## Supplementary Figures 1-5 for "Minimal activation of the p53 DNA damage response by a modular cytosine base editor enables effective multiplexed gene knockout in induced pluripotent stem cells"

#### SUPPLEMENTARY DATA

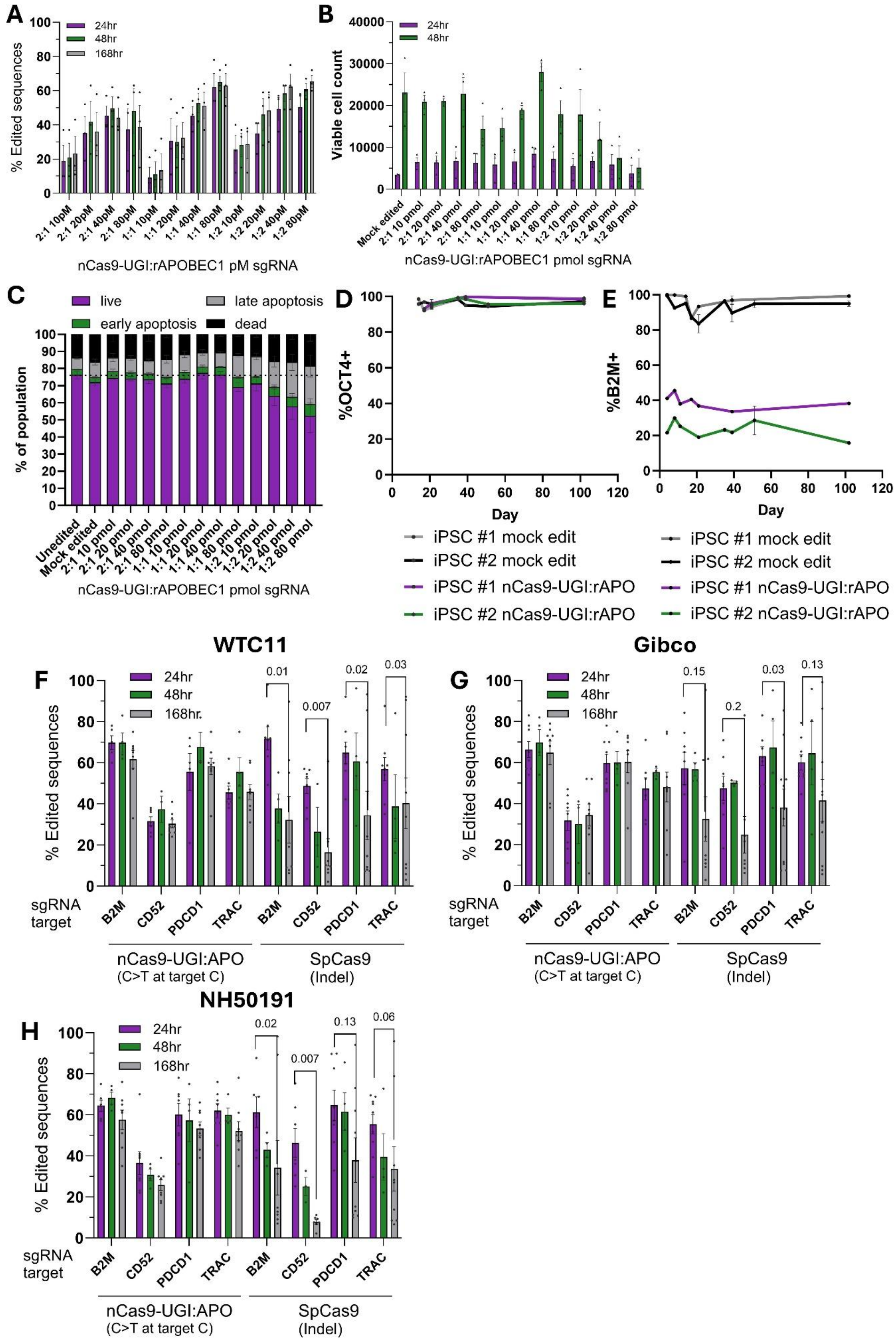

**Supplementary Figure 1:** Optimisation of efficient durable editing in iPSCs with an nCas9-UGI:rAPO base editor. A) Percentage of nCas9-UGI:APO-induced base conversion at the target nucleotide over time, B) Viable cell counts, and C) percentage of apoptotic cells following delivery of editor formulations composed of different molar ratios of nCas9-UGI and rAPOBEC-MCP mRNA and quantities of sgRNA while keeping the quantity of nCas9-UGI mRNA constant. Base conversion analysed by Sanger sequencing. Mean +/- SEM, n=3 iPSC lines for A-C. Percentage of cells expressing D) pluripotency marker OCT4 or E) target of knockout by base editing B2M during prolonged culture in edited pools at the indicated time following either mock editing or editing with the optimised nCas9-UGI:rAPO base editor formulation measured by flow cytometry.  Percentage of either SpCas9-induced indels or nCas9-UGI:APO-induced base conversion at the target nucleotide following delivery of editors targeting four genes simultaneously to the F) WTC11, G) Gibco, and H) NH50191 iPSC lines. Target base conversion or indel formation analysed at the indicated time points by Sanger sequencing. Mean +/- SEM, n=4-9. p-values from paired t-tests.

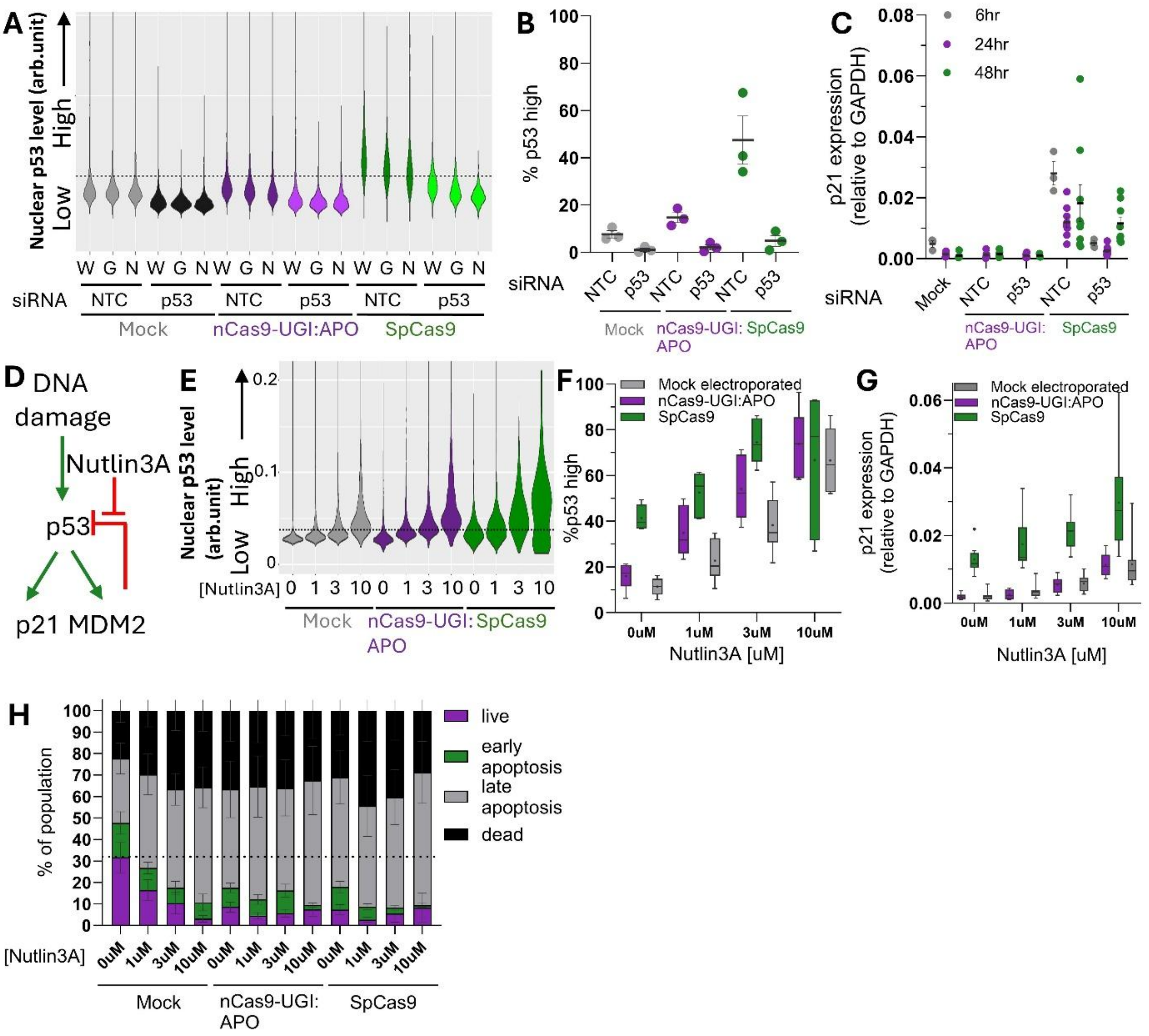


**Supplementary Figure 2:** Modulation of p53 pathway activation during multiplexed gene editing. A) Distributions of average nuclear p53 levels across three iPSC lines ((W)TC11, (G)ibco, (N)H50191) following multiplexed editing with either nCas9-UGI:APO or SpCas9 editors in the presence of either non-targeting control (NTC) or p53 targeting siRNA. p53 levels above the main distribution of mock edited controls were classified as 'high'. B) Percentage of p53 high nuclei following multiplexed editing with either nCas9-UGI:APO or SpCas9 in the presence of either non-targeting control (NTC) or p53 targeting siRNA. n = 3 iPSC lines. Mean +/- SEM. C) p21 expression following multiplexed editing with either nCas9-UGI:APO or SpCas9 editors in the presence of either non-targeting control (NTC) or p53 targeting siRNA. Analysed at the indicated timepoints following delivery of the editor. Mean +/- SEM, n=3-9. D) Schematic representation of de-repression of p53 levels by Nutlin3A inhibition of MDM2. E) Representative distributions of average nuclear p53 levels, F) percentage of p53 high nuclei (Tukey plots with mean shown by *, n= 6), and G) p21 expression (Tukey plots with mean shown by *, n=6-9), 24hrs following single target editing with either nCas9-UGI:APO or SpCas9 editors in the presence of the indicated concentrations of Nutlin3A. Quantification of H) Annexin V apoptosis assay. Mean +/- SEM, n=6.

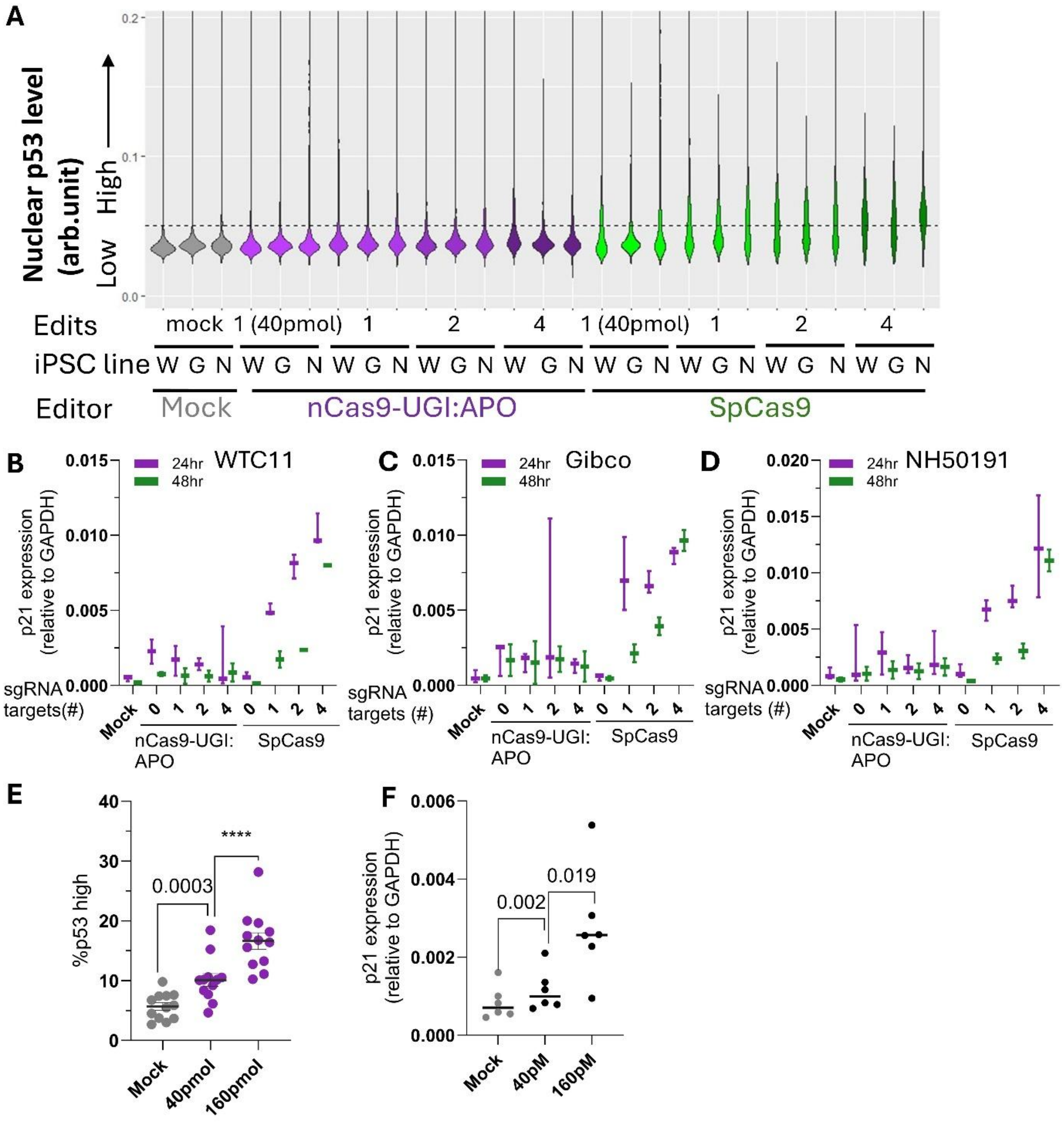


**Supplementary Figure 3**: p53 pathway activation by multiplexed gene editing. A) Distributions of average nuclear p53 levels across three iPSC lines ((W)TC11, (G)ibco, (N)H50191) following editing of the indicated number of gene targets with either nCas9-UGI:APO or SpCas9 editors. Total sgRNA quantity is 160 pmol unless indicated. p53 levels above the main distribution of mock edited controls were classified as 'high'. p21 expression following delivery of nCas9-UGI:APO or SpCas9 editors targeting the indicated number of genes simultaneously to the B) WTC11, C) Gibco, and D) NH50191 iPSC lines. Total sgRNA quantity was maintained at 160 pmol by addition of non-targeting control (NTC) sgRNA. Analysed at the indicated timepoints following delivery of the editor. Tukey plots with mean shown by *, n=3. Percentage of p53 high nuclei E), and p21 expression F), 24hrs following delivery of nCas9-UGI:APO with the indicated quantity of non-targeting control (NTC) sgRNA. Mean +/- SEM, n=12 for E and n=6 for F. p-values from paired t-tests; **** = p <0.0001.

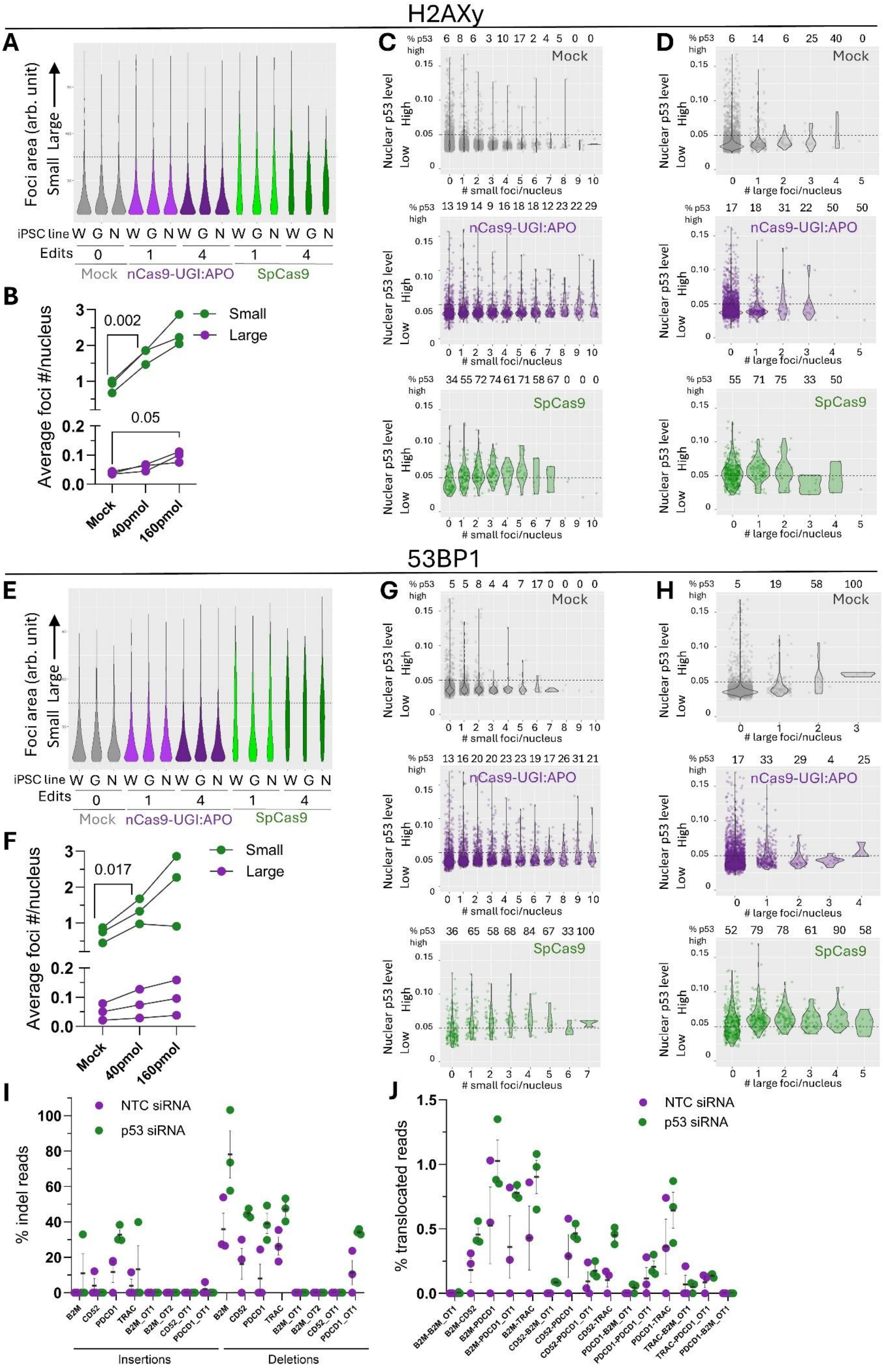
H2AXy
A
Foci area (arb. unit)
Small Large
iPSC line W G N W G N W G N W G N W G N
Edits 0 1 4 1 4
Mock nCas9-UGI:APO SpCas9
B
Average foci #/nucleus
0.002
0.05
Small
Large
Mock 40pmol 160pmol
C
% p53 high
6 8 6 3 10 17 2 4 5 0 0
Mock
Nuclear p53 level
Low High
### small foci/nucleus
13 19 14 9 16 18 18 12 23 22 29
nCas9-UGI:APO
34 55 72 74 61 71 58 67 0 0 0
SpCas9
D
% p53 high
6 14 6 25 40 0
Mock
### large foci/nucleus
17 18 31 22 50 50
nCas9-UGI:APO
55 71 75 33 50
SpCas9
53BP1
E
Foci area (arb. unit)
Small Large
iPSC line W G N W G N W G N W G N W G N
Edits 0 1 4 1 4
Mock nCas9-UGI:APO SpCas9
F
Average foci #/nucleus
0.017
Small
Large
Mock 40pmol 160pmol
G
% p53 high
5 5 8 4 4 7 17 0 0 0 0
Mock
Nuclear p53 level
Low High
### small foci/nucleus
13 16 20 20 23 23 19 17 26 31 21
nCas9-UGI:APO
36 65 58 68 84 67 33 100
SpCas9
H
% p53 high
5 19 58 100
Mock
### large foci/nucleus
17 33 29 4 25
nCas9-UGI:APO
52 79 78 61 90 58
SpCas9
I
% indel reads
NTC siRNA
p53 siRNA
B2M CD52 PDCD1 TRAC B2M_OT1 B2M_OT2 CD52_OT1 PDCD1_OT1 B2M CD52 PDCD1 TRAC B2M_OT1 B2M_OT2 CD52_OT1 PDCD1_OT1
Insertions
Deletions
J
% translocated reads
NTC siRNA
p53 siRNA
B2M-B2M_OT1 B2M-CD52 B2M-PDCD1 B2M-PDCD1_OT1 B2M-TRAC CD52-B2M_OT1 CD52-PDCD1 CD52-PDCD1_OT1 CD52-TRAC PDCD1-B2M_OT1 PDCD1-PDCD1_OT1 PDCD1-TRAC TRAC-B2M_OT1 TRAC-PDCD1_OT1 PDCD1-B2M_OT1

**Supplementary Figure 4:** Minimal NHEJ-mediated genome rearrangement associated with nCas9-UGI:rAPO base editing. A) Distributions of nuclear H2AXy foci sizes across three iPSC lines ((W)TC11, (G)ibco, (N)H50191) following editing of the indicated number of gene targets with either nCas9-UGI:APO or SpCas9 editors. H2AXy foci sizes above the main distribution of mock edited controls were classified as 'large'. B) Quantification of the average number of H2AXy foci per nucleus classified as either large or small 24hrs following delivery of nCas9-UGI:APO with the indicated quantity of non-targeting control (NTC) sgRNA. Mean +/- SEM, p-values from paired t-tests. Connecting lines show paired measurements from n=3 iPSC lines. Distributions of average nuclear p53 levels in pooled populations of nuclei from three iPSC lines containing the indicated number of either C) small, or D) large H2AXy foci. Analysed 24hrs following delivery of editors targeting 4 genes simultaneously. E) Distributions of nuclear 53BP1 foci sizes across three iPSC lines ((W)TC11, (G)ibco, (N)H50191) following editing of the indicated number of gene targets with either nCas9-UGI:APO or SpCas9 editors. 53BP1 foci sizes above the main distribution of mock edited controls were classified as 'large'. F) Quantification of the average number of 53BP1 foci per nucleus classified as either large or small 24hrs following delivery of nCas9-UGI:APO with the indicated quantity of non-targeting control (NTC) sgRNA. Mean +/- SEM, p-values from paired t-tests. Connecting lines show paired measurements from n=3 iPSC lines. Distributions of average nuclear p53 levels in pooled populations of nuclei from three iPSC lines containing the indicated number of either G) small, or H) large 53BP1 foci. Analysed 24hrs following delivery of editors targeting 4 genes simultaneously. Percentage of reads containing either I) insertions or deletions at all on target and known off-target sites, or J) translocations between on target and known off-target sites identified in one or more of n=3 iPSC samples edited with SpCas9 targeting four genes simultaneously in combination with either a non-targeting (NTC) or p53 targeting siRNA. Analysed by targeted sequencing 7 days following delivery of editors. Mean +/- SEM. n=3 iPSC lines.

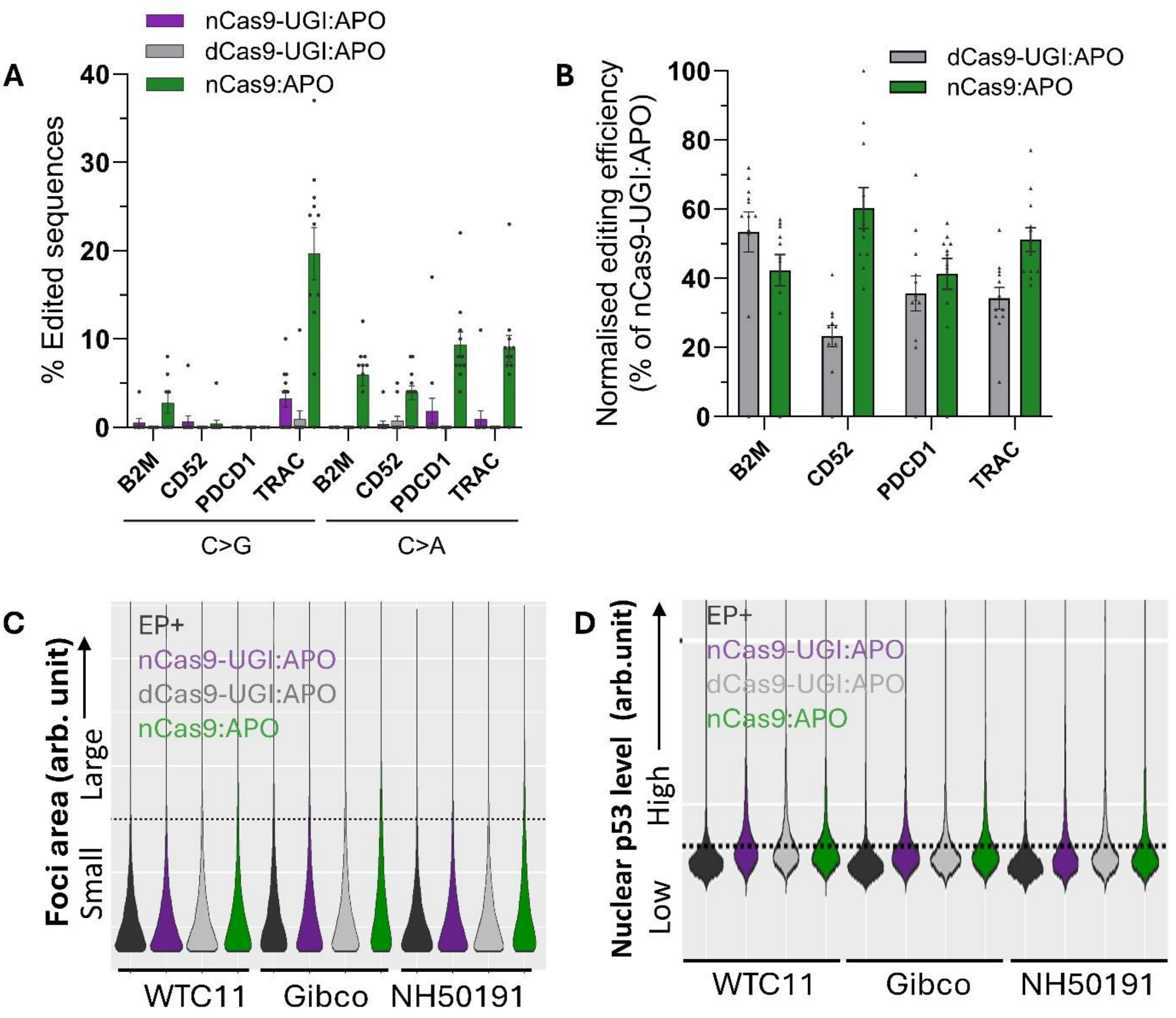


**Supplementary Figure 5:** Modulation of base-editor assembly minimises DSB formation by nCas9-UGI:rAPO. A) Percentage of C to non-T base conversions at the target nucleotides of four genes edited simultaneously following delivery of base editors composed of APOBEC1 and the indicated engineered SpCas9 variants. Editing analysed by sanger sequencing 7 days following delivery of editors. Mean +/- SEM, n=9-12. B) Percentage of base conversion at the target nucleotides of four genes edited simultaneously following delivery of base editors composed of APOBEC1 and the indicated engineered SpCas9 variants. Editing analysed by sanger sequencing 7 days following delivery of editors. Normalised editing efficiency expressed relative to matched control sample edited with nCas9-UGI:rAPO. Mean +/- SEM, n=12. Distributions of nuclear C) H2AXy foci sizes, and D) p53 levels, across three iPSC lines 24hrs following multiplexed editing of four target genes with base editors composed of APOBEC1 and the indicated engineered SpCas9 variants. H2AXy foci sizes and average nuclear p53 levels above the main distribution of mock edited controls were classified as 'large'.
